# Aberrant cortical reorganization and astrogliosis underlie memory deficit post-traumatic brain injury: a longitudinal multiscale multimodal study

**DOI:** 10.64898/2026.10.16.759571

**Authors:** Jazlynn Tan Xiu Min, Shruti Patel, Tony Xu, Adrienne Dorr, Dustin Loren V Almanza, Jonas Osmann, Thallita Rabelo, Laura Marie Vecchio, JoAnne McLaurin, Chao Wang, Nir Lipsman, Mathew Burke, Isabelle Aubert, Clement Hamani, Bojana Stefanovic, Maged Goubran

**Author notes:** Co-senior authors (Bojana Stefanovic, Maged Goubran).

## Abstract

We now have imaging and analytical tools to study the brain at multiple scales, from scanning the whole brain to see how different neuronal networks interact to scrutinizing individual cells. However, multimodal and multiscale investigations are challenging to conduct in the same subject and generate copious amounts of complex data. Such experimental designs are invaluable in studying heterogeneous neurological disease with pathology at both cellular and network level. Using traumatic brain injury as a test case, we conducted macroscale MRI imaging as well as 3D whole-brain light-sheet fluorescence microscopy with cellular resolution to study network dysfunction and markers of cognitive deficits post-injury. We identified sensitive neurophysiological markers that distinguish healthy and injured mice, and predicted their memory capabilities. The framework we present is broadly applicable to understanding complex cognitive functions and identifying the physiological correlates of distributed network dysfunction in neurological disorders.

**HIGHLIGHTS:**

● Multiscale framework integration of *in vivo* MRI and *ex vivo* whole-brain histology
● Connectomic gradients reveal network dedifferentiation linked to memory deficit
● 3D whole-brain segmentation show neuronal hyperactivity and widespread astrogliosis
● AI-based axon tracing reveals persistent damage despite normal neuroimaging scans

**SUMMARY:** Integrating high-dimensional neuroimaging data across spatial scales remains a fundamental bottleneck in quantitative neuroscience. Here, we apply emerging analytical techniques including functional connectome gradient mapping and 3D segmentation of whole-brain cellular resolution microscopy data in a mouse model of traumatic brain injury. At the macroscale, longitudinal functional MRI revealed a transition from frontal hyperconnectivity to hippocampal hypoconnectivity and unimodal-transmodal gradient contraction indicating network dedifferentiation. At the cellular scale, deep-learning based segmentation of whole-brain light-sheet fluorescence microscopy volumes quantified regional astrogliosis, neuronal hyperactivity, and single-axon loss undetected by standard diffusion MRI. Multiscale multimodal data integration not only identified quantitative biomarkers predictive of memory performance post-injury but also enabled validation of *in vivo* modalities against more sensitive *ex vivo* histology. This study demonstrates a framework for probing complex multi-network cognitive functions and neurological disorders of network dysfunction.

## INTRODUCTION

Advances in neuroimaging and fluorescence microscopy generate massive, high-dimensional datasets spanning orders of magnitude in spatial resolution and structural scale. Non-invasive magnetic resonance imaging (MRI) probes whole-brain functional and structural networks while light-sheet fluorescence microscopy (LSFM) enables rapid, cellular-resolution imaging of large brain tissue. While these modalities offer an unprecedented multiscale perspective of brain organization, extracting meaningful biological insights from these complex, high-dimensional data presents a major analytical bottleneck. At the macroscale, conventional resting-state functional MRI (rs-fMRI) analyses relied on discrete, pairwise regional correlations, failing to capture the continuous, topological organization of brain networks. This limitation is increasingly overcome by graph theory and dimensionality-reduction approaches like connectome gradient mapping. At the cellular scale, the difficulty of 3D segmentation of tera-voxel images often forces studies to sacrifice whole-brain approaches in favor of slice-based conventional histology of chosen region-of-interests (ROIs). Given recent advancement in deep-learning computer vision pipelines, we can now automate 3D segmentation and fiber tracing across intact whole-brain volumes.

Experimental mouse model of traumatic brain injury (TBI) is a good context in which to demonstrate the value of multi-scale, cross-modality tools. In TBI, mechanical forces shear axons that support communication within and between brain networks. Over weeks to months, neuroinflammation and metabolic disruption remodel the surviving circuitry^1–7^. Because injuries varying in force, location and severity converge on chronic memory impairment^8–12^, TBI is best understood as a distributed network disturbance rather than as focal damage^13^. It therefore warrants a whole-brain approach at both network and cellular scale. Clinically, TBI monitoring relies on non-invasive imaging that frequently lacks the sensitivity to reveal abnormalities particularly in milder injuries^14,15^. This underscores the need to validate macroscale signatures against sensitive cellular measurements. Marcoscale rs-fMRI investigations have demonstrated that functional reorganization post-TBI manifests as early global hyperconnectivity^16^, most pronounced in parietal and salience networks^17^, which settles into pericontusional hypoconnectivity^18^ and distal hyperconnectivity^19–21^. However, few such studies have associated macroscale findings with cellular level changes or performed histological validation. On the other hand, cellular scale studies primarily evaluated localized pathology only at chosen regions of interest (ROIs) often without linking observations to cognitive performance: subacute glial cell activation in the ipsi-contusional cortex^22^, global reduced neuronal activity^23^ and axon damage in the corpus callosum^24–26^.

In this study, we apply various analyses to *in vivo* MRI and *ex vivo* LSFM data acquired longitudinally in a repeated mild controlled cortical impact (CCI) model^27–29^ of TBI in mice to showcase how neuroscience can benefit from multiscale and crossmodality approaches. At the macroscale, we employed functional connectome gradients, which arrange regions along the continuous connectivity axes of brain organization rather than discrete pairs^30^. Changes to network position along connectomic gradients is sensitive to cognitive deficits but has not been applied to TBI^31–35^. We found subacute peri-contusional hyperconnectivity precedes hippocampal disconnection and a multi-network loss of functional specialization. At the cellular scale, we used tissue clearing and AI-based computational pipelines for analysis of whole-brain 3D LSFM data^36–38^. Astrogliosis describes the complex cellular response of astrocytes (regulators of neuroinflammation and synaptic plasticity^39–41^) to TBI^42^. LSFM revealed that astrogliosis, together with neuronal hyperactivity, correlated with memory impairment. Structurally, axonal tracing revealed persistent loss of injury site projections despite only transient diffusion MRI abnormalities. This work provides a scalable, reproducible computational approach for bridging macroscale network dynamics with whole-brain cellular histology to decode complex biological systems.

## RESULTS

### Mild injury results in chronic spatial memory impairment

We first established that our model reproduces the defining central challenge of mild TBI: an injury that does not leave overt damage yet still results in chronic memory impairment for weeks. In a pilot cohort, the repeated closed-head CCI paradigm lengthened the loss of righting reflex — the time a mouse takes to return from a supine to a prone position, a standard index of injury severity — relative to shams (sham: mean 21±0.5s; CCI: mean 202± 131s), with no apnea (Figures S1A and S1B). This range falls within what other studies classify as mild (<30s apnea, <6min loss of righting reflex, no skull fracture, no major hemorrhage)^43–46^. Consistent with a mild injury, we did not observe structural abnormalities on T2-weighted images (Figures S1D and S1E) and brain gray matter volumes exhibited no significant differences (Figure S1C).

Detecting the subtle deficits expected from such an injury demands a sensitive, clinically translatable memory test. We therefore assessed spatial memory using the location discrimination (LD) task in Bussey– Saksida touchscreen operant chambers (Figure 1B), in which a mouse learns to tell apart two nearby locations on a screen. Unlike conventional rodent paradigms such as the Morris water maze, the touchscreen platform is low-stress, closely parallels clinical assessments, such as the Cambridge Neuropsychological Test Automated Battery (CANTAB)^47,48^, and is more sensitive to subtle cognitive dysfunction^49,50^. On a high difficulty variant of the task (Figure 1B), sham mice improved their accuracy significantly from before to after surgery, as expected for normal learning, whereas CCI mice tested 7 weeks after injury failed to improve at all (shams: mean = +5.84, SD = 4.73, d = 0.564, *p <* 0.001; CCI: mean = - 3.19, SD = 8.38, d = -0.420, *p =* 0.377; paired permutation t-test, n = 5 per group with 5000 bootstrap samples) (Figure 1D). Having confirmed a chronic memory deficit on a clinically analogous task, we set out to trace its origin through the brain’s functional networks.

**Figure 1:**
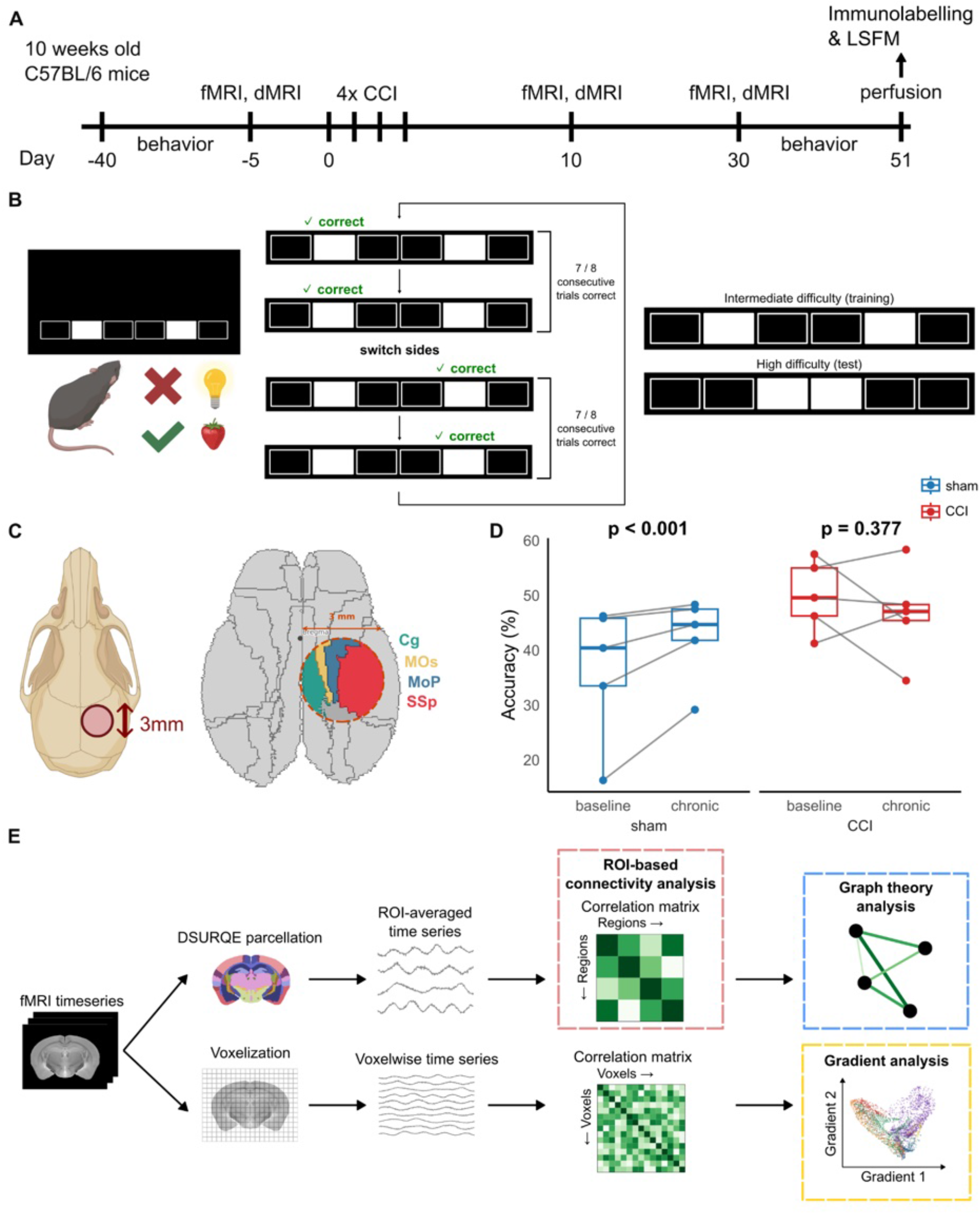
Experimental design and chronic cognitive impairment. (A) Ten-week-old C57BL/6 mice (sham: n=5, CCI: n=5) underwent training and testing for the LD task in touchscreen operant chambers. rs-fMRI and dMRI scans to assess functional and structural connectivity respectively were conducted 5 days prior to the four CCI hits or sham surgeries that were spaced by 24h. At the subacute and chronic timepoints – 10 and 30 days after the first CCI/sham, respectively – mice underwent rs-fMRI and dMRI scans. Performance on the LD task was reassessed 7 weeks post-injury and brain tissue was collected via transcardial perfusion. SHIELD-based clearing and immunolabelling were conducted on whole-brain tissue, which was then imaged using light-sheet fluorescence microscopy. (B) In the touchscreen operant chamber, the mouse must select the correct (left or right) lit square to receive a strawberry milkshake reward; an incorrect selection triggers an aversive light. When 7 out of 8 consecutive correct selections were made, the correct side switched. Mice were trained on intermediate difficulty (right, top) and tested on high difficulty (right, bottom). (C) The location of the 3-mm diameter CCI (-1.5 AP, 1.5ML) on the right hemisphere over the skull (left) and the key regions of interest relative to the impact site (right). (D) The CCI group failed to improve in accuracy at the chronic stage compared to baseline, while the sham group showed a significant increase in accuracy (permutation t-test, 5000 permutations). (E) Analysis framework for rs-fMRI.

### Functional connectivity disruption migrates from the injury site to the hippocampus over time

To locate where communication breaks down, we used resting-state functional MRI (rs-fMRI), which images spontaneous brain activity while the animal is at rest. From these scans, we measured functional connectivity: the degree to which the activity of two regions rises and falls together over time. For each pair of ROIs, connectivity was defined as the Pearson correlation between their mean BOLD signals over a 45-min scan (Figure 1E). Group-averaged correlation matrices, normalized by Fisher’s z-transformation, are shown in Figure S2A. An initial exploratory survey of the whole connectivity matrix showed a transient bilateral increase in connectivity within the injury site (motor, parietal and cingulate cortices) at the subacute stage. Reduced connectivity within the hippocampal formation, by contrast, appeared only chronically (Figure 2A).

**Figure 2:**
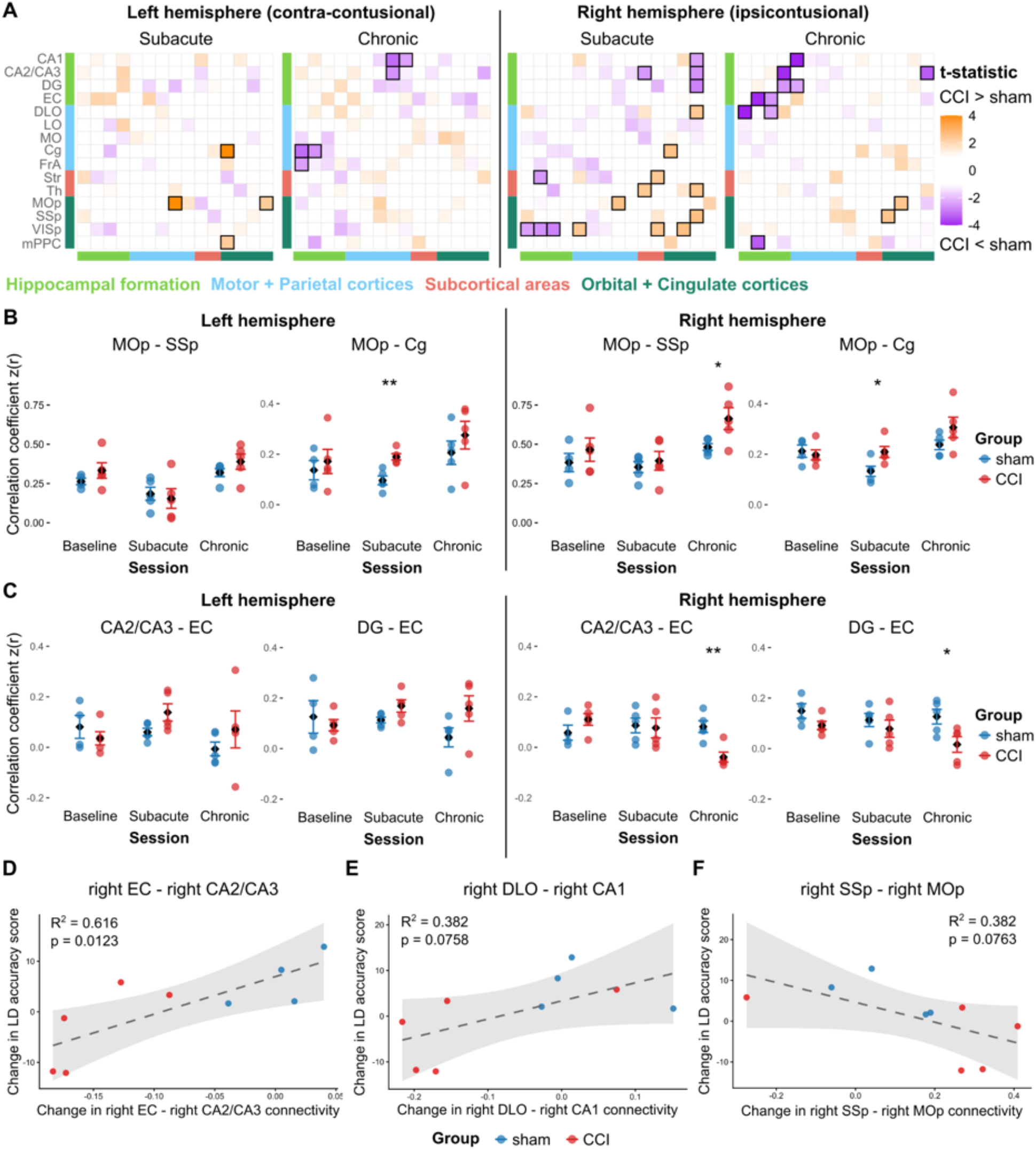
rs-fMRI revealed transient pericontusional hyperconnectivity and chronic hippocampal hypoconnectivity. (A) Groupwise comparisons of z-scored correlations between pairs of ROIs within the same hemisphere at the subacute and chronic timepoints. Color indicates the t-statistic of regression, where a positive t-statistic indicates that the CCI group (n=5) had a higher z-scored correlation than the sham (n=5) group. A negative t-statistic is where the CCI group had a smaller z-scored correlation than the sham group. Significant (*p <* 0.05) t-statistics are highlighted with a thick border. (B) Functional connectivity between the Cg and the MOp (injury site) was bilaterally increased transiently at the subacute stage in the CCI group. At the chronic stage, functional connectivity between the MOp and SSp at the injury site was increased ipsi-contusionally in the CCI group. (C) Functional connectivity between regions of the ipsi-contusional hippocampal formation (CA2/CA3 - EC and DG - EC) was decreased chronically in the CCI group. No significant differences were observed contra-contusionally. (D) Improvement in LD accuracy score was positively correlated with increased right EC - CA2/CA3 connectivity. (E) A similar trend was observed for increased right DLO - right CA1 connectivity (see Figure S2B for groupwise comparisons), and (F) a negative trend with increased right SSp - right MOp connectivity. LD accuracy score improvement and connectivity change were calculated as the difference between the chronic and baseline readouts. In (B) and (C), points represent individual animals colored by group, diamonds indicate group means and error bars represent standard error. In (D) - (F), each point represents one animal, colored by group. The dashed line shows the ordinary least-squares regression fit, where the shaded band indicates the 95% confidence interval of the regression line. See Supplementary Table 1 for abbreviations. Asterisks indicate p-value, where *p <* 0.05 *, *p <* 0.01 **, *p <* 0.001 ***.

We tested this pattern more rigorously in an a priori set of ROIs at the CCI site and within the learning-memory circuits (Supplementary Table 1). Subacutely, connectivity between the primary motor cortex ‘MOp’ to cingulate ‘Cg’ in the injury site was elevated in both hemispheres (left: t = 4.30, d = 2.72, *p <* 0.01, *p_corr_* < 0.01; right: t = 2.58, d = 1.63, *p <* 0.05, *p_corr_* < 0.1) (Figure 2B). By the chronic stage, this pericontusional hyperconnectivity was only observed on the ipsi-contusional side between the MOp and primary somatosensory cortex ‘SSp’ (t = 2.48, d = 1.57, *p <* 0.05, *p_corr_* = 0.114) (Figure 2B). The hippocampus followed the opposite time course: no significant change subacutely, but chronic diminished connectivity on the injured side between CA2/CA3 and entorhinal cortex (t = −3.93, d = −2.48, *p <* 0.01, *p_corr_* < 0.05) and between the dentate gyrus and entorhinal cortex (t = −2.59, d = −1.64, *p <* 0.05, *p_corr_* < 0.1) (Figure 2C). The injured brain thus follows a biphasic, spatially migrating course: an early, recoverable surge of connectivity at the impact site progressed to chronic disconnection of the hippocampus which is a key hub in the memory network.

Critically, these connectivity changes were associated with memory performance across animals. Greater improvement in LD accuracy was associated with a larger increase in ipsi-contusional CA2/CA3–entorhinal cortex connectivity from baseline (R^2^ = 0.616, f^2^ = 1.60, *p <* 0.05, *p_perm_* < 0.05, 95% CI R^2^: [0.114, 0.918]) (Figure 2D). Two further relationships trended in the same direction: better memory improvement accompanied a greater increase in ipsi-contusional dorsolateral orbital cortex–CA1 connectivity (R^2^ = 0.382, f^2^ = 0.619, *p =* 0.0758, *p_perm_* = 0.0709, 95% CI R^2^: [0.023 0.745]) (Figure 2E) and a larger decrease in ipsi- contusional SSp–MOp connectivity (R^2^ = 0.382, f^2^ = 0.617, *p =* 0.0763, *p_perm_* = 0.0719, 95% CI R^2^: [0.0300, 0.630]) (Figure 2F). Preserved hippocampal connectivity thus accompanied better recovery, whereas lingering perilesional hyperconnectivity accompanied memory impairment.

### Brain networks become more segregated in the chronic phase post-CCI

We next asked whether these pericontusional and hippocampal connectivity changes reorganized the whole-brain network or remained localized. We used graph theory to investigate whether injury disturbs the balance between functional segregation and integration (Figure 1E). Each animal’s network was summarized with three metrics: clustering coefficient (how tightly a region’s neighbors interconnect, i.e. local segregation), local efficiency (how well information flows within a local neighborhood), and global efficiency (how efficiently information travels across the whole network, i.e. integration). To avoid arbitrary thresholding bias^51^, we evaluated each metric across a range of network densities and tested whether this density dependence differed between CCI and sham groups.

Subacutely, no measure differed between groups across densities. At the chronic stage, relative to sham, the CCI network had lower global efficiency (Figure 3A) and a higher clustering coefficient (Figure 3C) (global efficiency: t = 3.69, d = 0.458, *p <* 0.001, *p_corr_* < 0.001; clustering coefficient: t = −2.71, d = -0.336, *p <* 0.01, *p_corr_* = 0.108), with the difference most pronounced in sparse networks of the strongest connections. Local efficiency was indistinguishable between groups (Figure 3E). For closer examination, we fixed the threshold at a density similar to prior work^16,31,52,53^(top quartile of connections, 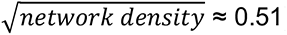), and emphasize the strong connections that are disproportionately important for information flow across the network (Figures 3A and 3C). At this density, the CCI group again showed lower global efficiency (b = - 0.0247, t = −2.50, d = −1.58, *p <* 0.05, *p_corr_* = 0.111) (Figure 3B and 3D) and a trend toward higher clustering (b = 0.0658, t = 2.04, d = 1.29, *p =* 0.0763, *p_corr_* = 0.114). Together, these results indicate that by the chronic stage the injured brain had become more segregated (higher clustering) yet less integrated (lower global efficiency). The network had fragmented into locally clustered groups while losing some of the long-range links that normally bind them together.

**Figure 3:**
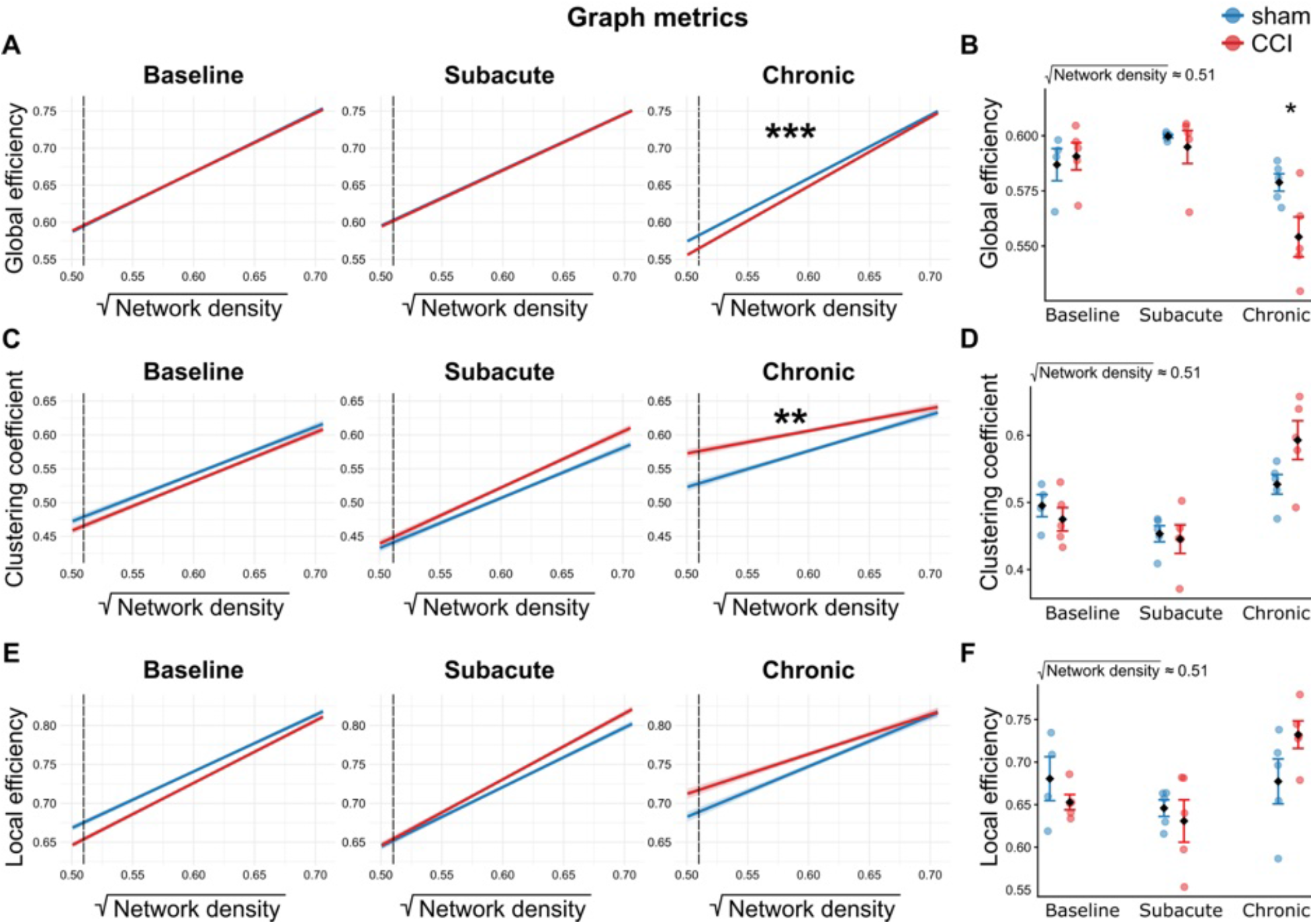
Graph-theory analysis highlights network segregation at the chronic stage. Global efficiency (A) was lower and clustering coefficient (C) higher in CCI (n=5) relative to sham (n=5) across network densities, with the group difference increasing toward lower densities (significant group × network-density interaction); mean local efficiency (E) showed no such interaction. The dashed line represents the density with the strongest quartile of connections (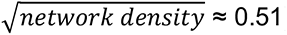). At this density, the CCI group had lower global efficiency (B) and higher clustering coefficient (D), while mean local efficiency did not differ (F). In (A), (C) and (E), lines represent model-predicted means while shaded regions indicate ± standard error of the fitted values. In (B), (D) and (F), points represent individual animals colored by group, black diamonds indicate group means, and error bars represent standard error. Asterisks indicate *p*-value, where *p <* 0.05 *, *p <* 0.01 **, *p <* 0.001 ***.

### CCI contracts the brain’s functional hierarchy

As parcellated brain atlases cannot capture the continuous hierarchy of functional organization, we mapped each voxel along axes of "connectome gradients" based on functional connectivity similarity^30,31^ (Figure 1E). Each animal’s gradients were aligned to a common template built from the group-averaged baseline connectome. Voxels were then grouped into functional networks as described by prior work^31^. The first gradient captured an executive-to-mnemonic axis within the transmodal cortex, and the second separated transmodal from unimodal (primary sensory) regions (Figure 4A), reproducing the mouse functional gradients reported previously^31,54^ and together explaining 31% of the variance (Figure 4B). Left hemisphere functional connectivity gradients are shown in Figure S3B. The hippocampal network of CCI mice expanded along the executive–mnemonic axis relative to shams (t = −1.99, d = −1.26, *p =* 0.081) (Figures 4C and 3D) — a spreading-apart of hippocampal voxels that echoes the within-hippocampus disconnection seen in ROI-based connectivity (section 2.2). At the same time, the frontal and auditory networks contracted toward the center of the gradient space, drifting away from the transmodal and unimodal poles, respectively (frontal: t = −3.37, d = −2.13, *p <* 0.01, *p_corr_* < 0.1; auditory: t = 2.51, d = 1.59, *p <* 0.05, *p_corr_* = 0.108) (Figures 4E and 3F). The magnitude of this contraction of the ipsi-contusional frontal network was associated with LD accuracy (R^2^ = 0.756, f^2^ = 3.09, *p <* 0.01, p_perm_ < 0.01, 95% CI R^2^: [0.153, 0.971]) (Figure 4G) — a stronger correlate of memory performance than any individual ROI-based connectivity.

**Figure 4:**
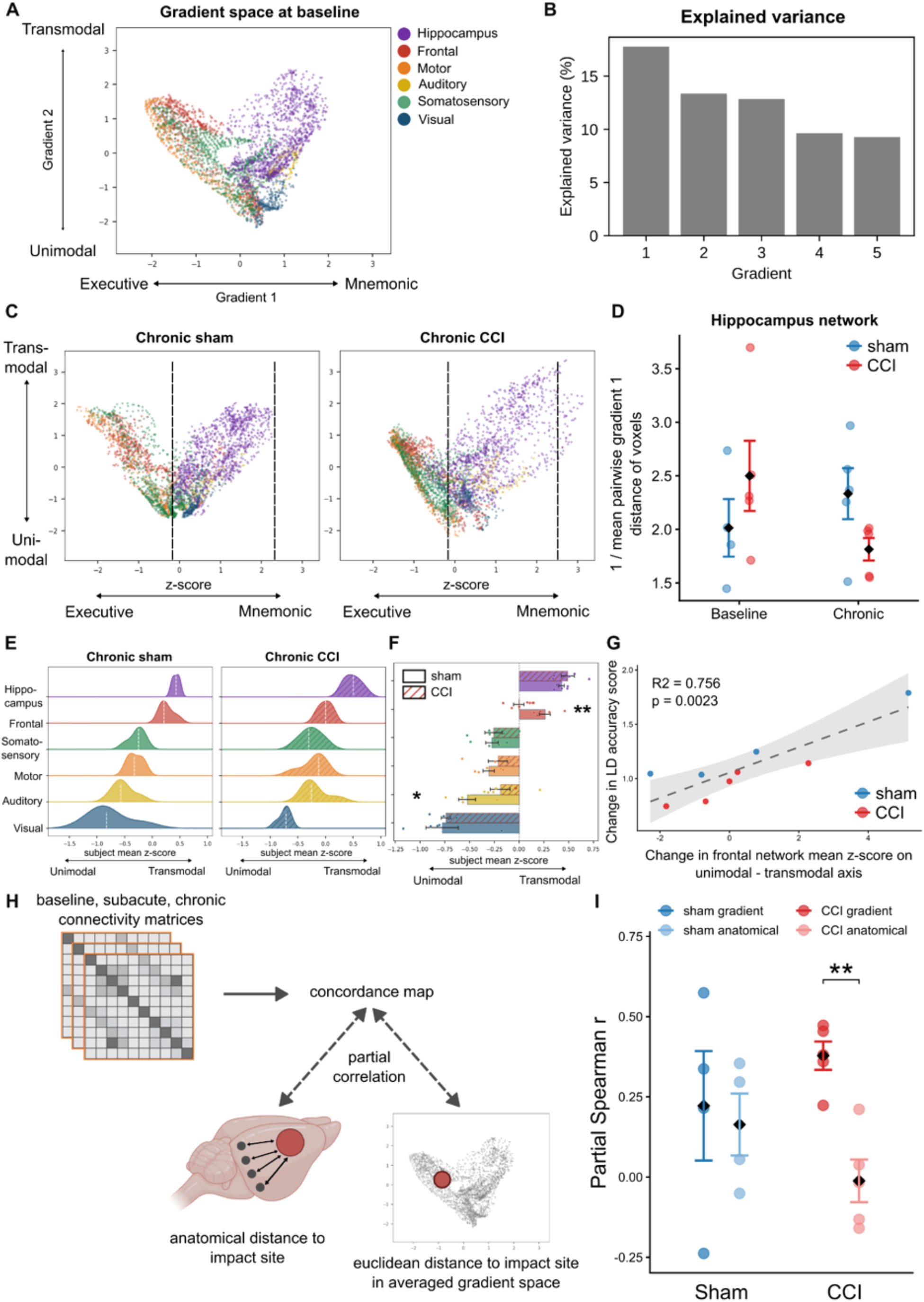
Functional connectome gradients contract post-TBI. (A) Baseline template gradient, defined by gradient 1: transmodal executive - transmodal mnemonic and Gradient 2: unimodal - transmodal. Each point represents a voxel in the fMRI, colored by the functional network it belongs to. See Figure S3A for group and session template gradients. (B) Gradient 1 and 2 explain 31% of the variance. Gradient space visualization (C) and quantification as the reciprocal of the mean pairwise distance among voxels in the network (D) showed chronic decrease in hippocampal compactness in CCI versus shams (n=5 per group). (E) Distribution of z-scored network centroids along Gradient 2 (mean value per network; dashed lines indicate medians). (F) At the chronic phase, there was a significant contraction of the frontal network from the transmodal pole and the auditory network from the unimodal pole in the CCI group. (shaded bars indicate CCI; each point represents the network centroid of a subject. (G) The improvement in LD accuracy score was significantly positively correlated with the shift of the frontal network centroid. Change was calculated as chronic readout divided by baseline readout. (H) For each subject, the voxelwise temporal concordance matrix was correlated with the Euclidean distance from the impact site in gradient and physical space. (I) The partial Spearman r of gradient distance was significantly larger than that of anatomical distance in the CCI group, but not in the sham group. In (D) and (I), points represent individual animals colored by group, diamonds indicate group means and error bars represent standard error. In (G), each point represents one animal, colored by group. The dashed line shows the ordinary least-squares regression fit, where the shaded band indicates the 95% confidence interval of the regression line. Asterisks indicate *p*-value, where *p <* 0.05 *, *p <* 0.01 **, *p <* 0.001 ***.

Prior work had shown that connectivity changes propagated with gradient-space rather than anatomical distance to the stroke lesion^32^. We thus asked whether the longitudinal temporal concordance of each voxel’s connectivity profile was better explained by its distance to the contusion in gradient or physical space. We used partial correlations to isolate each predictor’s unique contribution while controlling for their spatial correlation (Figure 4H). In CCI animals, gradient-space distance was a strong and reliable predictor of concordance independent of physical distance, with a positive partial correlation in all five animals (mean r = 0.378; one-sample t = 7.92, *p <* 0.01, d = 3.54) (Figure 4I). In contrast, anatomical distance carried essentially no unique information (mean r = −0.012; *p =* 0.872). The per-animal difference confirmed that gradient distance contributed more than anatomical distance (mean difference = 0.390; t = 5.13, *p <* 0.01, d = 2.29). In shams, neither predictor reached significance, and the two did not differ (mean gradient r = 0.222; mean anatomical r = 0.164; difference *p =* 0.790). A blanket variance-attenuation effect would instead suppress both equally, implicating what we observed as injury-specific reorganization rather than a floor effect. Permutation tests, via shuffling concordance across voxels, collapsed the partial r to near zero in every CCI animal confirming the connectome gradient effect (Fisher combined *p* < 0.001; Figure S3D). Together, these results indicate that the loss of longitudinal connectivity stability after TBI is organized by a region’s position on the functional connectivity manifold rather than by its physical proximity to the lesion, consistent with disruption propagating along connectivity gradients.

### Contra-contusional hyperactivity post-CCI is associated with memory impairment

Resting-state connectivity is a proxy of the brain’s intrinsic functional architecture at the macroscopic level, but some deficits may emerge only when that architecture is placed under demand. A circuit whose resting connectivity looks preserved may still fail to engage during memory retrieval, and the two measures can dissociate. To capture task-evoked activity at cellular resolution, we collected brain tissue 1h after the mice performed the intermediate difficulty LD task and immunolabeled for the immediate-early gene c-Fos that labels recently active neurons (Figure 5A-D). The injured hemisphere was hypoactive (Figures 5E and 4F), with significantly fewer c-Fos+ cells in the subiculum, a major hippocampal output (t = −2.94, d = −1.86, *p <* 0.05, *p_corr_* < 0.1) (Figure 5G). The intact hemisphere was instead hyperactive, most prominently in the primary auditory cortex (t = 2.41, d = 1.53, *p <* 0.05, *p_corr_* < 0.1) (Figure 5H), likely driven by the task’s auditory feedback cues. This contra-contusional hyperactivity was maladaptive rather than compensatory: the more active the contra-contusional auditory cortex, the less the animal’s memory improved (R^2^ = 0.596, f^2^ = 1.48, *p <* 0.01, *p_perm_* < 0.05, 95% CI R^2^: [0.008, 0.898]) (Figure 5I), implicating sensory-network dysfunction in the memory deficit.

**Figure 5:**
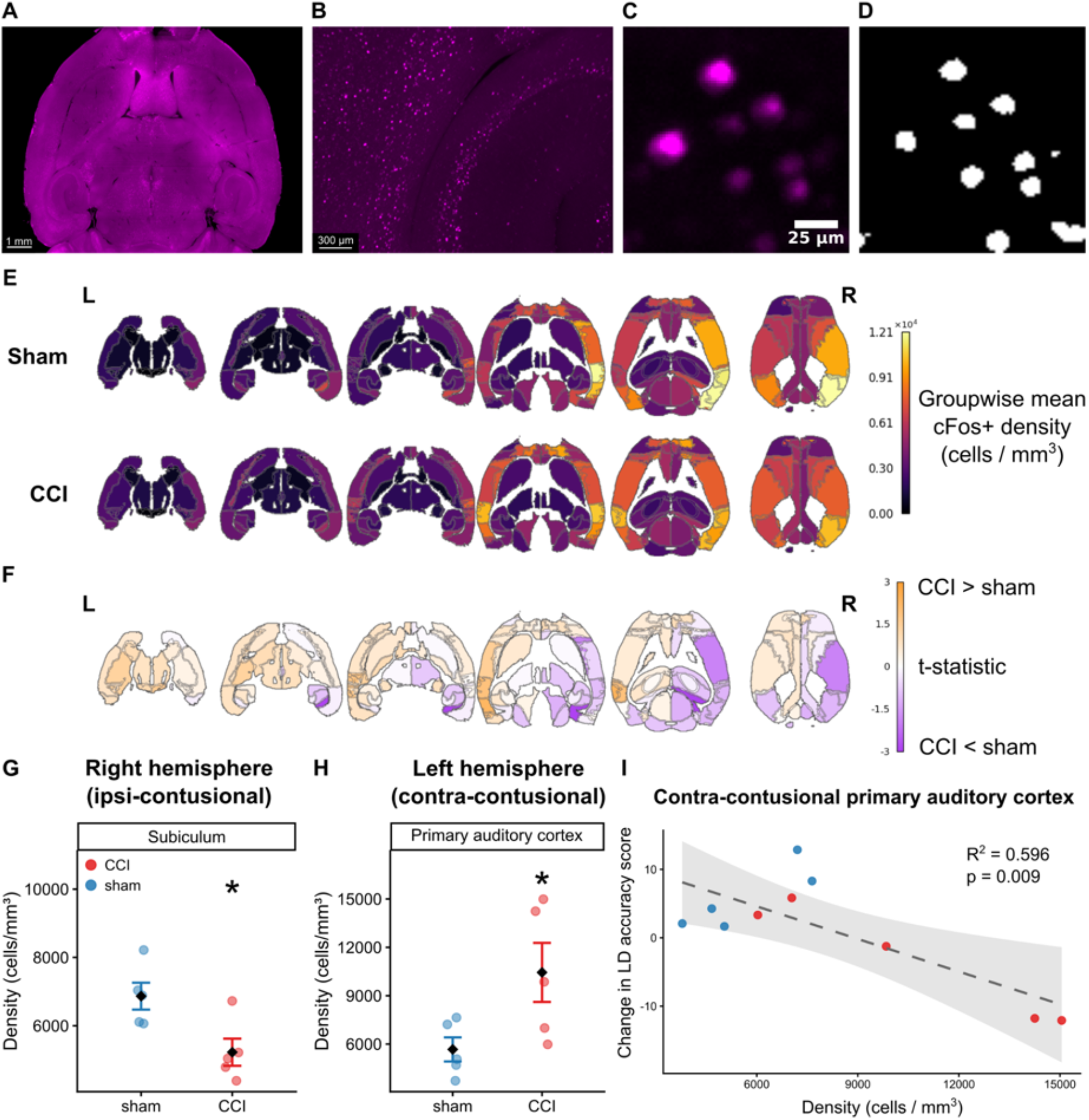
CCI resulted in ipsi-contusional hypoactivity and contra-contusional hyperactivity. (A) Maximum intensity projection (40 µm) of a LSFM whole-brain image of cFos staining, with a magnified view of the hippocampus (B). (C) Representative cFos+ cells and corresponding segmentation mask (D). (E) Group-averaged cFos+ cell density maps show a decrease ipsi-contusionally and an increase contra- contusionally in the CCI (n=5) group compared to shams (n=5). Color scale shows group-averaged cFos+ cell density. (F) T-statistics from region-wise groupwise comparisons. Color scale shows t-statistic. Regions in orange indicate higher cFos+ cell density in CCI than in shams, while regions in purple indicate lower cFos+ cell density in CCI than in shams. (G) cFos+ cell density decreased in the ipsi-contusional subiculum and (H) increased in the contra-contusional primary auditory cortex. (I) The increase in neuronal activation in the contra-contusional primary auditory cortex was significantly correlated with poorer improvement in the LD accuracy score from baseline to chronic timepoint. In (G) and (H), points represent individual animals colored by group, diamonds indicate group means, and error bars represent standard error. In (I), each point represents one animal, colored by group. The dashed line shows the ordinary least-squares regression fit, where the shaded band indicates the 95% confidence interval of the regression line. Asterisks indicate *p*-value, where *p <* 0.05 *, *p <* 0.01 **, *p <* 0.001 ***.

### Diffusion MRI detected transient loss of crossing fibers

We next sought to investigate the structural underpinnings of the functional reorganization. We used *in vivo* diffusion MRI (dMRI) to study white matter microstructure at both the subacute and chronic stages, focusing on four tracts chosen for their vulnerability to injury or their role in visuospatial memory: the corpus callosum, internal capsule, fornix, and fimbria. Subacutely, fractional anisotropy (FA) was elevated in CCI mice in the ipsi-contusional internal capsule (t = 2.34, d = 1.48, *p <* 0.05, *p_corr_* < 0.1) and fimbria (t = 2.79, d = 1.77, *p <* 0.05, *p_corr_* < 0.1) (Figure 6B), alongside reduced orientation dispersion (ODI; fimbria: t = −2.87, d = −1.81, *p <* 0.05, *p_corr_* < 0.1; internal capsule: t = −2.65, d = −1.67, *p <* 0.05, *p_corr_* < 0.1) and no change in neurite density (NDI) (Figures 6D and 5F). To determine whether these effects extended beyond our four a priori tracts, we performed an exploratory whole-brain analysis spanning both hemispheres (Figures 6G–I). No tract in the left hemisphere showed a group difference in FA, NDI, or ODI, indicating that the microstructural changes were confined to the right ipsi-contusional hemisphere (Figures 6A, 5C and 5E). This signature of higher FA and lower ODI suggests a selective loss of crossing fibers in these tracts. By the chronic stage, these changes were no longer detected. We also found no change in the corpus callosum or fornix at either timepoint, despite their well-documented vulnerability in human TBI^55,56^.

**Figure 6:**
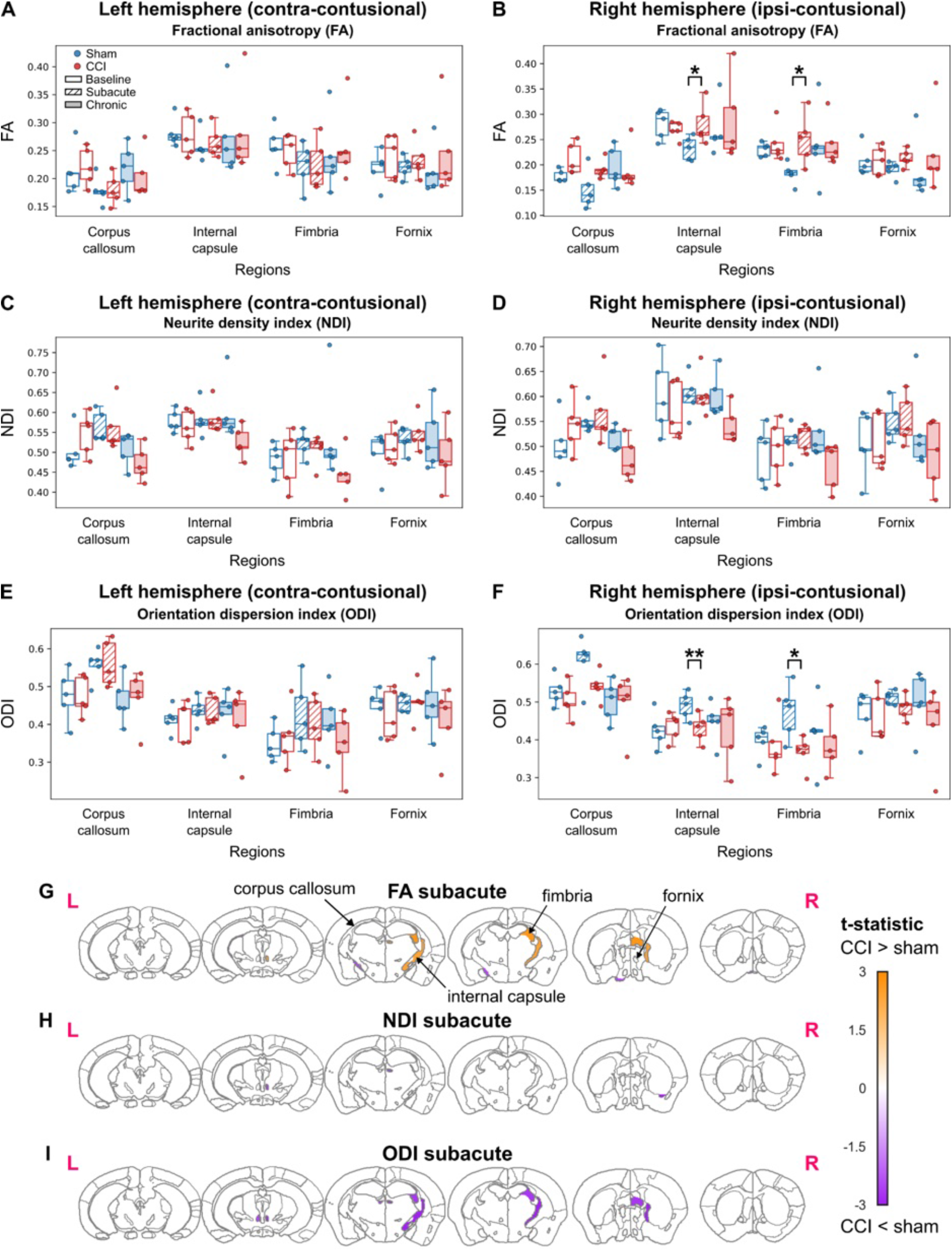
dMRI reveals white matter changes subacutely after CCI. Fractional anisotropy (FA) in the (A) left contra-contusional and (B) right ipsi-contusional hemispheres. FA was significantly higher in the CCI (n=5) group compared to shams (n=5) in the right ipsi-contusional internal capsule and fimbria at the subacute stage. No significant differences in neurite density index (NDI) were found in the same regions in either the contra-contusional (C) or ipsi-contusional (D) hemispheres. (E) Orientation dispersion index (ODI) in the (E) contra-contusional and (F) ipsi-contusional hemispheres. Significant decreases were detected in ODI in the ipsi-contusional internal capsule and fimbria at the subacute stage. Exploratory whole-brain analysis for FA (G), NDI (H), and ODI (I) at the subacute stage, where all white matter regions with significant t-statistics (*p <* 0.05) are shown. Color indicates t-statistic from an ordinary least squares model modeling the group effect at the subacute or chronic sessions. Orange regions had larger metric values in the CCI group compared to the sham group. Purple regions had smaller metric values in the CCI group compared to the sham group. Asterisks indicate *p*-value, where *p <* 0.05 *, *p <* 0.01 **, *p <* 0.001 ***.

### Single-axon tracing reveals persistent, asymmetric loss of projections

*in vivo* dMRI has low spatial resolution, which limits our analysis to large fiber bundles, and diffusion metrics lack microstructural specificity. To test whether axonal damage resolves at the single-axon level, we injected an anterograde viral tracer into the cortex under the center of the impact. We then imaged labeled axons in the cleared whole brain and mapped them in 3D using MAPL3^37^ (Figures 7A-C). Unlike the apparent recovery on dMRI, axonal projection densities were chronically reduced brain-wide in CCI mice (Figures 7D and 6E). The steepest decreases were on the side opposite the injury in the fimbria (t = -4.43, d = −2.97, *p <* 0.01, *p_corr_* < 0.01), superior colliculus (t = -4.27, d = −2.87, *p <* 0.01, *p_corr_* < 0.05), and ventral orbital cortex (t = −3.06, d = −2.06, *p <* 0.05, *p_corr_* < 0.05), marking degeneration of cross-hemispheric projections (Figure 7F). On the injured side, projections into the hippocampus were also diminished, significantly in the dentate gyrus (t = −2.52, d = −1.69, *p <* 0.05, *p_corr_* < 0.1) and as a trend in the fimbria (t = −2.00, d = −1.34, *p =* 0.085, *p_corr_* = 0.136) (Figure 7G). Because raw axon density depends on tracer uptake for each animal, we also computed a lateralization index (LI = (right - left) / (right + left)) that is normalized within each animal. This showed that degeneration was uneven across hemispheres: the dentate gyrus had a lower LI in CCI mice, reflecting disproportionate loss on the injured side (t = −3.09, d = −2.07, *p <* 0.05), whereas the striatum (t = 2.52, d = 1.69, *p <* 0.05) and insular/ectorhinal cortex (t = 2.31, d = 1.55, *p =* 0.054) had a higher LI (Figures 7H and 6I), reflecting the selective vulnerability of transcallosal corticostriatal and cortico-cortical projections^25,57–59^. We can hence infer that the structural injury did not recover, but instead likely fell below the detection threshold of *in vivo* dMRI.

**Figure 7:**
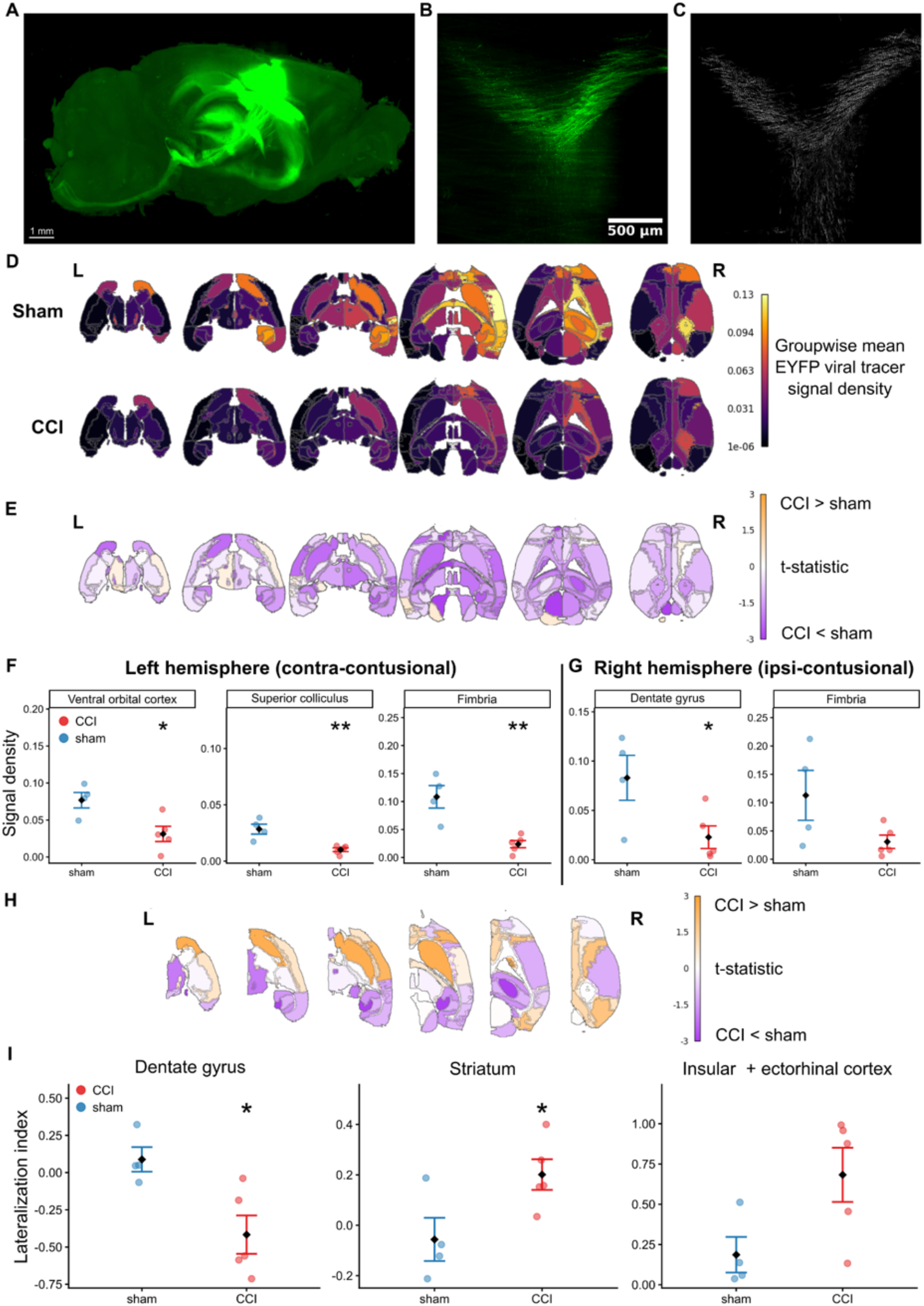
CCI induced cross-hemispheric and cortico-hippocampal axonal loss. (A) Maximum intensity projection (40 µm) of a LSFM whole-brain image of EYFP labelled axons, with a magnified view of the corpus callosum (B) and corresponding segmentation mask (C) (D) Group-averaged signal density maps show a global decrease in the CCI (n=5) group compared to shams (n=5). Color scale shows group-averaged signal density from segmented axons masks. (E) T-statistics from region-wise groupwise comparisons. Color scale indicates t-statistic where regions in orange indicate higher EYFP signal density in CCI than in shams; while regions in purple indicate lower EYFP signal density in CCI than shams. (F) Significant decreases in EYFP signal density in contra-contusional ventral orbital cortex, superior colliculus and fimbria; and (G) ipsi-contusional dentate gyrus and fimbria. (H) T-statistics from region-wise groupwise comparisons of within-hemisphere normalized lateralization index (LI). Color scale indicates t-statistic where regions in orange indicate higher LI in CCI than shams while regions in purple indicate lower LI in CCI than shams. (I) LI was decreased in dentate gyrus, but elevated in the striatum and insular + ectorhinal cortex regions in the CCI group. In (F), (G), and (I), points represent individual animals colored by group, diamonds indicate group means and error bars represent standard error. Asterisks indicate *p*-value, where *p <* 0.05 *, *p <* 0.01 **, *p <* 0.001 ***.

### Ipsi-contusional astrocytic reactivity persists chronically

Besides axon shearing, astrogliosis is a major component of secondary TBI injury. Furthermore, there is growing interest in the role of astrocytes in learning and memory. To map this response brain-wide, we immunolabeled cleared tissue for GFAP at the subacute and chronic stages (Figures 8A-D). Before comparing groups, we normalized GFAP signal to whole-brain levels to correct for differences in staining efficiency and for the baseline inflammation that the tracer injection produced in both groups (Figures 8E and 7F).

**Figure 8:**
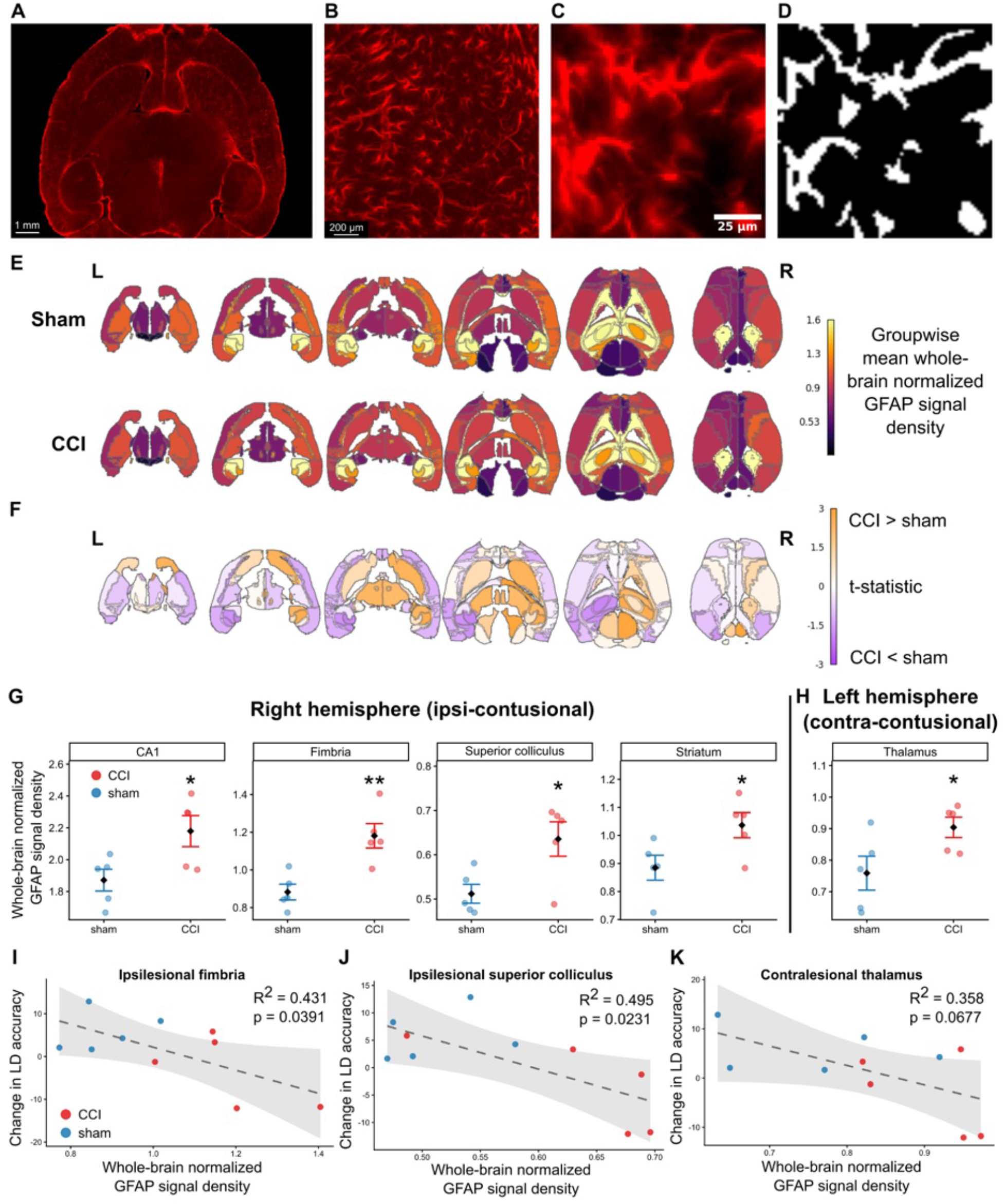
CCI results in chronic ipsi-contusional astrogliosis. (A) Maximum intensity projection (40 µm) of a LSFM whole-brain image of GFAP staining, with a magnified view of the hippocampus (B). (C) Representative GFAP+ cells and corresponding segmentation mask (D). (E) Group-averaged whole-brain normalized GFAP signal density maps show a global decrease in the CCI (n=5) group compared to shams (n=5). Color scale shows group-averaged whole-brain normalized signal density from segmented GFAP+ astrocytes. (F) T-statistics from region-wise groupwise comparisons. Color scale indicates t-statistic, where regions in orange indicate higher GFAP signal density in CCI than shams while regions in purple indicate lower GFAP signal density in CCI than shams. (G) Whole-brain normalized GFAP signal density increased in ipsi-contusional CA1, fimbria, superior colliculus, and striatum in the CCI group. (H) Whole-brain normalized GFAP signal density increased in contra-contusional thalamus in the CCI group. Whole-brain normalized GFAP signal density in the ipsi-contusional fimbria (I) and superior colliculus (J), and in the contra-contusional thalamus (K), was negatively correlated with improvement in LD accuracy score from baseline to chronic timepoint. In (G) and (H), points represent individual animals colored by group, diamonds indicate group means, and error bars represent standard error. In (I) - (K), each point represents one animal, colored by group. The dashed line shows the ordinary least-squares regression fit, where the shaded band indicates the 95% confidence interval of the regression line. Asterisks indicate p-value, where p < 0.05 *, p < 0.01 **, p < 0.001 ***.

In a small pilot cohort (n=3) and exploratory analysis, subacute GFAP reactivity was concentrated near the injury site, in primary motor and somatosensory cortex (Figure S4A). By the chronic stage, this perilesional reactivity had partly subsided, even as GFAP across the wider ipsi-contusional hemisphere appeared broadly elevated relative to subacute CCI mice (Figure S4B).

At the chronic stage, normalized GFAP was significantly elevated in CCI mice on the injured side in the CA1 (t = 2.58, d = 1.63, *p <* 0.05), fimbria (t = 3.87, d = 2.45, *p <* 0.01, *p_corr_* < 0.05), superior colliculus (t = 2.79, d = 1.77, *p <* 0.05, *p_corr_* < 0.1), and striatum (t = 2.40, d = 1.52, *p <* 0.05, *p_corr_* < 0.1) (Figure 8G), regions that notably overlap the sites of axon loss above, linking the inflammation to the degenerating projections. As with the axon data, a lateralization index captured the hemispheric distribution: LI was increased in the corpus callosum (t = 2.60, d = 1.64, *p <* 0.05, *p_corr_* < 0.05), CA1 (t = 3.18, d = 2.01, *p <* 0.05, *p_corr_* < 0.05), and secondary visual cortex (t = 2.56, d = 1.62, *p <* 0.05, *p_corr_* < 0.05) (Figures S4C and S4D), driven by lower contralateral signal alongside preserved or slightly raised ipsi-contusional signal. Memory improvement was negatively correlated with this chronic GFAP in the ipsi-contusional fimbria (R² = 0.431, f^2^ = 0.758, *p <* 0.05, *p_perm_* < 0.05, 95% CI R^2^: [0.007, 0.795]) (Figure 8I) and superior colliculus (R² = 0.495, f^2^ = 0.982, *p <* 0.05, *p_perm_* < 0.05, 95% CI R^2^: [0.030, 0.800]) (Figure 8J). In the contra-contusional thalamus, GFAP was elevated in CCI mice (t = 2.31, d = 1.46, *p =* 0.050) (Figure 8H) and trended toward predicting worse memory (R² = 0.358, f^2^ = 0.557, *p =* 0.068, *p_perm_* = 0.0669, 95% CI R^2^: [0.013, 0.811]) (Figure 8K), echoing the burden of ipsilateral astrogliosis on memory performance.

## DISCUSSION

Although the brain continues to reorganize at the cellular and network level to affect memory performance post-injury, few studies have characterized these changes through a multiscale lens. We attribute this to three major challenges: the oversimplification of high-dimensional fMRI data into discrete, pairwise regional correlations; the computational complexity of segmenting and quantifying cellular pathology in teravoxel 3D brain volumes; and the scarcity of matched, cross-modal datasets that validate non-invasive clinical MRI against sensitive cellular histology in the same animal. To address these gaps, we demonstrate an integrative framework combining connectomic gradient analysis that represents functional brain organization on continuous axes of variation, deep-learning-based 3D segmentation of LSFM data, and within-subject validation of *in vivo* dMRI via viral axon tracing. At the macroscale, several fMRI analyses converged on chronic hippocampal hypoconnectivity and loss of network specialization as drivers of memory deficits. At the cellular scale, whole-brain LSFM segmentation revealed that astrogliosis, even in regions distal to the primary injury site, correlated with memory impairment. Finally, axonal loss persisted chronically despite the normalization of diffusion MRI metrics, highlighting the sensitivity limits of non-invasive clinical neuroimaging. Thus, the analytical framework we present provides a strategy for decoding multiscale network reorganizations in neurological disorders.

### Connectome gradient analysis reveals a compression of cortical hierarchy beyond conventional connectomics

In this study, conventional connectomics revealed chronic within-hippocampus hypoconnectivity, in agreement with the expansion of the hippocampal network in gradient space, as a loss of shared connectivity drives nodes apart. Crucially, functional gradient analysis uncovered chronic compression of functional hierarchy, where the auditory and frontal networks drifted inward from their respective unimodal and transmodal poles. Together with the chronic reduction in global graph efficiency, the hierarchical compression points toward a targeted breakdown of long-range functional pathways. While conventional pairwise ROI connectomics pinpoint localized aberrations, graph representations show network-level changes, and gradients capture the brain’s continuous functional organization.

Bayrak et al. demonstrated that in humans, post-stroke functional connectivity changes propagate along gradient axes rather than radiating anatomically outwards from the lesion^32^. Our data agree with these findings, where in injured animals, gradient distance correlated more strongly than anatomical distance with the amount of change in functional connectivity. Moreover, the most extensive chronic functional connectivity changes occurred in the hippocampal network, a region remote from the CCI site in the primary motor and somatosensory cortex. This suggests that the dysfunction spread through a region’s functional connections rather than its physical neighborhood. The gradient framework is also valuable for predicting cognitive outcomes. Shen et al. found that in humans, post-stroke memory impairment was associated with the de-differentiation of the frontoparietal-visual and frontoparietal-somatomotor network pairs^33^. In healthy individuals, several studies have also found that gradients predict cognition with similar or better accuracy than traditional parcellation-based connectivity^60,61^. Consistent with this, we found that the shift in the frontal network position along the unimodal-transmodal gradient axis predicted memory performance better than any of the ROI-ROI pairwise connectivity strengths.

### Whole-brain imaging overcomes sampling bias to reveal distal and heterogeneous pathology

Conventional 2D histology causes physical disruption of spatial continuity across cut sections and necessitates *a priori* region selection, as manually analyzing each slice is unfeasible. We instead combined intact whole-brain LSFM to preserve 3D structural integrity with deep learning-based segmentation to perform brainwide quantitative analysis. Applied to c-Fos mapping, we found a marked hemispheric asymmetry in memory task-evoked neuronal activity following CCI, with ipsi-contusional hypoactivity alongside contra-contusional hyperactivity. The reduction in activity in the ipsi-contusional subiculum agreed with prior reports of post-TBI hypoactivity: Frankowski et al. demonstrated large reductions in visually evoked potential amplitudes and firing rate after an open skull CCI^62^, and Anwer et al. reported a global decrease in c-Fos+ cell density 6h after a rotational head injury^23^. By extending neuronal activity mapping to a task-engaged context, we additionally identified hyperactivity in contra-contusional primary auditory cortex, a region engaged by the LD task’s auditory feedback cues, which correlated with poorer LD accuracy improvement. This may reflect maladaptive compensation, similar to how aberrant synaptogenesis and contra-contusional engagement are associated with poorer recovery post-stroke^63–65^. Additionally, preferential vulnerability of cortical inhibitory interneurons to CCI^62,66^ could drive transcallosal disinhibition, resulting in hyperactive but noisy contralateral signaling^67,68^. Our findings therefore position sensory network dysfunction as potentially integral, rather than peripheral, to the higher-order cognitive impairment that follows injury^69–72^.

Similarly, whole-brain GFAP mapping captured diffuse, heterogeneous astrogliosis that would have been overlooked by typical 2D sampling often restricted to the injury site. The pronounced ipsi-contusional astrocytic reactivity one week post-injury fits the secondary injury cascade, in which neuroinflammatory signaling propagates along anatomically connected circuits^73,74^. Yet, chronic-phase reactivity has been variably reported, with some studies describing resolution^75–77^ while others (including this study) demonstrating persistent activation^78,79^. Notably, TBI biases neural stem cells away from astrogenesis toward neurogenesis, leaving pre-existing mature astrocytes responsible for post-TBI GFAP elevation^80^. However, post-TBI neurogenesis is frequently dysfunctional, marked by aberrant dendritic branching and ectopic cellular migration^81–83^. Without sufficient concurrent astrocytic support to stabilize the local microenvironment and modulate synapses^40,84,85^, the influx of dysfunctional neurons can further undermine local circuits. The combination of 3D LSFM imaging and deep learning-based quantification presented here can support deeper investigations involving transcriptomic profiling to distinguish astrocyte subtypes^86–89^ for a more thorough understanding of astrogliosis post-TBI.

### Axonal tracing exposes subtle chronic microstructural changes undetected by dMRI

Using dMRI, we identified a transient loss of crossing fibers in the ipsi-contusional fimbria and internal capsule at the subacute stage, which appeared to resolve by the chronic timepoint. In contrast, high-resolution viral tract tracing revealed persistent reductions in projection density across the brain, particularly in the contra-contusional hemisphere. Notably, both modalities aligned in detecting no structural changes within the corpus callosum or fornix, supporting the notion that these specific interhemispheric tracts are spared by the reduced rotational shear characteristic of closed-head injury models^90–93^. Where the modalities diverged, however, the comparison underscores the limited sensitivity of *in vivo* dMRI.

We propose three mechanisms to explain why dMRI might have missed the persistent pathology. First, axonal loss or regrowth may have been too fine or dispersed to survive partial-volume averaging within dMRI’s large voxels, thus remaining below the detection threshold^94,95^. Other studies have likewise reported histological white matter pathology that was undetected by dMRI^96,97^. Second, dMRI infers microstructural properties from water diffusion hence metrics can appear unchanged if opposing effects offset one another. For example, the hypertrophy of fibrous astrocytes arranged parallel to axons in white matter tracts^98,99^ can raise diffusion anisotropy while axon loss counteracts it^100^, leading to normalization. Third, axons vary in both vulnerability and regenerative capacity according to their orientation, location, and neuronal subtype^58,101–104^. dMRI indiscriminately measures all white matter components, while the viral tracer selectively labels the CaMKII+ projection neurons from the injury site. The discrepancy may thus be partially explained by differences in the axonal population under scrutiny.

Our anterograde tract tracing complements prior work by Frankowski et al., which used retrograde rabies tracing from surviving somatostatin interneurons in the hippocampus and prefrontal cortex after focal CCI and reported reduced long-range inputs in mice^105^. Similarly, our lateralization analysis showed that the long-range cross-hemispheric excitatory projections to the striatum and cortex were among the most disproportionately damaged contra-contusionally. Together, these findings demonstrate that focal TBI triggers chronic structural remodeling that can be invisible on *in vivo* neuroimaging and highlights the need for validation of clinical scans against more sensitive measures.

### Limitations of the study

Sex differences in post-TBI neurophysiology and cognitive outcomes have been reported in humans and rodents^106–109^. We included a mix of male (N=4, 2 per group) and female (N=6, 3 per group) mice, but the small sample sizes precluded stratified analyses, requiring animals to be pooled across sexes. Additionally, while we conducted brain-wide analyses to fully leverage our macroscale modalities, these remain exploratory given the limited statistical power. Consequently, our primary analyses focused on an *a priori* selection of brain regions and functional networks directly implicated in the injury site and memory performance. A second limitation is the use of anaesthesia during rs-fMRI acquisition, which could reconfigure functional connectivity in dose- and agent-dependent ways^110,111^. Although our isoflurane– dexmedetomidine protocol is standard in rodent MRI, its interaction with the injured brain may confound results. Awake recordings during a memory task would be more sensitive to the neurophysiological consequences of injury and are a natural extension of this work. Finally, our repeated TBI model with 24h injury interval, while common amongst rodent studies^112–114^, re-injures the brain during its acute injury phase. In humans, repeat injuries typically when the preceding injury has entered its chronic phase. We used shorter intervals to account for accelerated pathobiology in mice^115^, ensure reliable cognitive deficits^116,117^ and avoid age-related confounders from a longer experimental timeline. Nevertheless, this temporal difference is an important consideration for clinical translation.

## Conclusion

This study presents a multiscale approach that bridges *in vivo* imaging with cellular resolution *ex vivo* histology to map the trajectory of network reorganisation following TBI. Connectomic gradient mapping revealed that post-injury dysfunction propagates along functional hierarchies rather than simple anatomical distance, leading to loss of network specialization that predicted injury-induced memory deficits. Furthermore, deep-learning based 3D segmentation of LSFM volumes showed widespread, chronic cellular pathology and exposed the sensitivity limits of standard clinical diffusion MRI in detecting subtle axonal damage. Ultimately, this work demonstrates how multimodal and multiscale investigations can deepen our understanding of complex neurological disorders and inform more targeted clinical management.

## RESOURCE AVAILABILITY

### Lead contact

Requests for further information and resources should be directed to and will be fulfilled by the lead contact, Maged Goubran.

### Materials availability

This study did not generate new unique reagents.

### Data and code availability

Behavioral, raw MRI and regionwise axon/cell density from microscopy data have been deposited at the Federated Research Data Repository as https://doi.org/10.20383/103.01833 and the Open Data Commons for Traumatic Brain Injury (ODC-TBI) as [to be available after acceptance] and are publicly available as of the date of publication. All original code has been deposited at https://github.com/AICONSlab/MouseCCI-MRI-LSFM-submission.git and is publicly available as of the date of publication.

## ACKNOWLEDGMENTS

This work was supported by funding from Canadian Institute of Health Research (CIHR) grants PJT178059, PJT376309, and PJT156179, National Institutes of Health (NIH) grant R01AG084681-01, Natural Sciences and Engineering Research Council (NSERC) Discovery grant RGPIN-2021-03728 and the Weston Brain Institute. J.T.X.M. is supported by the National Science Scholarship (PhD) awarded by the Agency for Science, Technology and Research (A*STAR), Singapore. C.H. was supported by a Tier 1 Canada Research Chair in Translational Research. B.S. is funded by the Canada Research Chairs program award #CRC-2018-00042. M.G. is supported by the Canada Research Chairs program award #CRC-2021-00374. We are grateful to the Digital Research Alliance of Canada (alliancecan.ca) for their allocation of computing resources used in parts of this research. We also thank Andrew Chen from Queen’s University for his help in annotating ground truth labels for GFAP+ cells in microscopy images.

## AUTHOR CONTRIBUTIONS

Conceptualization, J.T.X.M., B.S. and M.G.; Data collection, J.T.X.M., S.P., A.D., D.L.V.A., T.R. and L.M.V.; Data analysis, J.T.X.M. and T.X.; writing – original draft, J.T.X.M., B.S. and M.G; writing – review & editing, J.T.X.M., S.P., A.D., D.L.V.A., T.X., L.M.V., M.B., I.A., C.H., B.S. and M.G.; funding acquisition, I.A., B.S.

and M.G.; supervision, B.S. and M.G.

## DECLARATION OF INTERESTS

C.H. is a consultant for Abbott.

## DECLARATION OF GENERATIVE AI AND AI-ASSISTED TECHNOLOGIES IN THE WRITING PROCESS

During the preparation of this work, the author(s) used Claude and Gemini in order to streamline writing of analysis code and proof-reading of the manuscript. After using this tool/service, the author(s) reviewed and edited the content as needed and take(s) full responsibility for the content of the published article.

## FIGURE TITLES AND LEGENDS

Embedded in-text for initial submission.

## METHODS

### Animals and Surgeries

All experiments and procedures in this study adhered to ARRIVE 2.0 guidelines^118^ and were approved by the Animal Care Committee of Sunnybrook Research Institute, which adheres to the policies and guidelines of the Canadian Council on Animal Care and meets all requirements of the provincial Statute of Ontario, Animals for Research Act, and the Canadian Federal Health of Animals Act. Sixteen 10-week-old C57BL6 mice (Charles River Canada) were housed in a 12h light/dark cycle with free access to water and food except during behavioral training and testing. Figure 1A shows the experimental timeline. The repeated closed-skull controlled cortical impact (CCI) model involved four hits with a 24h inter-impact interval. During each CCI, mice were induced with 4% isoflurane and maintained with 2% isoflurane on a 70% medical air and 30% oxygen mix. 5mg/kg Baytril and 1ml of Lactated Ringer’s was administered subcutaneously and the mouse was maintained at a body temperature of 37°C on a feedback-controlled warming pad affixed to a stereotaxic frame. After administration of 0.8mg/kg marcaine at the scalp, the skull was exposed via a midline incision. The 3mm metal impact tip was centered over the right somatosensory cortex (1.5mm posterior to bregma and 1.5mm lateral to the midline) (Figures 1B and 1C) by tilting the mouse such that the tip was parallel to the skull surface. The impact was delivered via the Leica Impact One Stereotaxic Impactor at a velocity of 4 m/s, to a depth of 1mm and a dwell time of 200ms. For sham mice, the impactor tip was lightly touched to the skull without impact. A burr hole was then made at the impact site (1.5mm AP, 1.5mm ML) to allow 1µl of 1×10^13^ genome copies/ml AAV2/DJ8-CaMKIIa-EYFP-NRN (Canadian Neurophotonics Platform Viral Vector Core Facility, RRID:SCR_016477) to be injected 1 mm into the somatosensory cortex over 10 min through a glass micropipette. The needle was left in place for 3 min before retraction. The incision was sutured and the mouse allowed to recover in a heated cage. Four mice were used for a preliminary assessment of the CCI severity. Neurological scoring ranging from 0 (no deficit) to 3 (severe deficit) for forelimb flexion, hindlimb flexion, and limb placing was performed before and after CCIs. Immediately after the impact or light touch (for shams), the duration of apnea and loss of righting reflex were recorded. The remaining 12 mice underwent behavioral testing and *in vivo* MRI scans, as detailed in Figure 1A. Two mice died in the course of the MRI scanning and were excluded from subsequent analyses.

### Operant chamber behavior

To ensure motivation, mice were lightly restricted to 90-95% free-feeding weight while accounting for normal growth during behavioral testing. Mice were trained to perform the location discrimination (LD) task in the Bussey-Saksida touchscreen operant chamber. Training took place daily during the light cycle (08:30– 12:30), after which they were weighed and fed. The mice learned to use the operant chamber through a multi-stage process^49,119,120^, before training specifically for LD. During training, 2 boxes at intermediate distance (boxes 2 and 5 out of 6 boxes arranged horizontally equal distances apart) lit up on the screen. Mice had to nose-poke the correct side to receive a strawberry milkshake reward while an incorrect touch triggered an aversive light punishment (Figure 1D). After touching the correct side 7 out of 8 times consecutively, the correct side reversed. 50 trials were allowed per session, and mice were considered to have successfully learned the task after completing at least 2 reversals each for 2 consecutive sessions. Following this, mice underwent testing for 4 challenging LDsmall sessions (smaller distance between boxes – boxes 3 and 4 lit) (Figure 1D) and 4 easy LDlarge sessions (larger distance between boxes – boxes 1 and 6 lit).

### MRI acquisition

Mice were scanned using a 7T horizontal bore MRI scanner (BioSpec 70/30 USR with BGA-12SHP gradients running ParaVision 6.0.1, Bruker BioSpin, Billerica, MA, USA). Face clamps secured the mouse head in position, and a 20-mm diameter loop receive coil was placed over the animal’s head while an 86- mm diameter volume coil was used for excitation. Mice were induced with 4% and maintained on 2% isoflurane on a 70% medical air and 30% oxygen mix during positioning of the mouse. After ensuring the brain was in the isocenter, a bolus of 0.05mg/ml dexmedetomidine diluted in saline at a concentration of 0.255mg/kg body weight was delivered subcutaneously. After 3 minutes, isoflurane was reduced by 0.25% every minute until a final concentration of 0.25%. Ten minutes after bolus delivery, a subcutaneous infusion of dexmedetomidine at 0.255mg/kg/hr was started. Breath rate was monitored via a respiration pillow sensor and maintained at approximately 90–110 breaths per min by increasing or reducing isoflurane within the 0–0.5% range as needed. One hour after infusion, dexmedetomidine infusion rate was lowered to 0.191 mg/kg/hr to maintain stable physiology. Throughout the scan, the mice were kept warm with a heated water blanket.

The T2-weighted structural scan was acquired coronally via a rapid acquisition with refocused echoes (RARE) sequence (TR/TE = 2500/40.89ms, slice thickness = 1mm, FOV = 19.2x19.2mm, in-plane resolution=0.15x0.15mm, pixel bandwidth = 256.99 Hz, flip angle = 90°). Map shimming was run to minimize B0 inhomogeneities. The resting state functional MRI (rs-fMRI) scan was acquired coronally using a spin echo echo planar imaging (SE-EPI) sequence (TR/TE = 2000/30.05ms, slice thickness = 1mm, FOV = 19.2x19.2mm, in-plane resolution = 0.30x0.30mm, pixel bandwidth = 3125 Hz, flip angle = 90°) for 45 minutes. Diffusion-weighted scans (dMRI) were acquired with a multi-shell protocol with EPI readout (TR/TE = 4500/29.54ms, slice thickness 1mm, FOV = 19.2x19.2mm, in-plane resolution=0.30x0.30mm, pixel bandwidth =3125 Hz, flip angle = 90°, b values = 0, 1215 and 1928s/mm^2^, with 33 equally spaced directions). An additional scan in the opposite encoding direction was acquired for b-value = 0 for EPI distortion correction.

### Brain volume analysis

T2-weighted structural scans were preprocessed via a method adapted from Pagani et al.^121^. Images were first upsampled to 0.090 mm isotropic voxel size, then bias corrected using *N4BiasFieldCorrection* (shrink factor = 2, spline spacing = 180, spline order = 3, convergence levels = 50) in the ANTs toolkit. The DSURQE atlas^122–125^ image was registered to the corrected images through rigid, affine, and symmetric image normalization registration using *antsRegistrationSyNQuick*. Skull stripping was carried out using the mask generated from the union of the output from FSL’s *bet* and the warped DSURQE atlas mask. A second round of bias correction and registration to DSURQE atlas space was performed on the skull-stripped images. The DSURQE atlas labels were transformed into anatomical native space.

### Functional connectivity

rs-fMRI scans were first denoised using *DenoiseImage* from the ANTs toolkit and all volumes were registered to the mean across the volumes for each scan. Denoising and a rigid registration improved the quality of preprocessing steps in RABIES^126^. Preprocessing comprised inhomogeneity correction of anatomical and functional scans, slice timing correction, and registration to the DSURQE template space. 24 motion parameters^127^ and timecourses from the first 5 principal components derived from voxels in white matter and cerebrospinal fluid^128^ were modeled at each voxel and regressed from the data using ordinary least-square regression. Finally, signal variability was homogenized to prevent confounding high-variance voxels from dominating ROI averages used in connectivity estimation. To preserve statistical power, we modified the parcellations of the DSURQE atlas to reduce the number of ROIs and limited subsequent analyses to these. Finally, connectivity matrices were generated by computing Pearson’s r between ROI pairs.

### Graph theory

For graph theory analyses, we thresholded the connectivity matrix to retain the top 20–50% of connections at 1% gradations prior to binarization. This reduces biases stemming from a predetermined threshold^16^. Lower densities emphasize the strongest connections, while higher densities include weaker ones. Using *bctpy*, we calculated three network metrics: (1) global efficiency: the average inverse shortest path length, indicating how readily information flows across the entire network; (2) mean local efficiency: the average inverse shortest path length among a node’s neighbors after that node is removed, indicating information flow within local neighborhoods; and (3) mean clustering coefficient: the mean, across nodes, of the ratio of actual to possible connections among a node’s neighbors, quantifying local interconnectedness. We first fit a regression model with an interaction term between group and 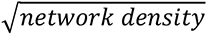, which shows whether group differences in these network metrics vary across connection strength. Then, after identifying the network threshold where there was greatest difference between groupwise means, we performed sham vs. CCI comparisons via a separate regression model, without the interaction term.

### Functional connectome gradients

A voxelwise connectivity matrix was constructed from voxels within the cortex and hippocampus of each registered rs-fMRI scan. The matrix was thresholded to retain the top 20% of connections and Fisher-transformed. Group-averaged connectomes were created by taking the mean across each group at each timepoint. Then, five principal gradients were extracted from both the group-averaged connectomes and individual scan connectomes using diffusion embedding and cosine kernel in *brainspace*. For the right hemisphere, we analyzed the first and second gradients which explained a combined 31% of variance. For the left hemisphere, we analyzed the first and third gradients which explained a combined 29% of variance. All group-averaged and individual scan gradients were aligned to the sham group’s baseline, through Procrustes alignment with 100 iterations and a tolerance level of 1×10^-6^. Within-subject z-score normalization was performed for each gradient. The mean gradient value of all voxels in each network in each gradient was compared between groups. Network compactness was defined as the reciprocal of the mean pairwise distance between all pairs of voxels within the network.

Using the same voxelwise connectivity matrix, we computed a voxelwise longitudinal concordance map to quantify the stability of every voxel’s functional connectivity profile across the baseline, subacute, and chronic sessions^129^. Two competing predictors of concordance were defined relative to the CCI contusion ROI: (i) gradient-space distance, the Euclidean distance of each voxel to the contusion centroid in the first two connectivity gradients, averaged across the analysis sessions; and (ii) anatomical distance, the physical Euclidean distance (mm) of each voxel to the contusion centroid in world space^32^. The contusion ROI was defined from the CCI target in the right hemisphere up to 0.8mm into the cortex based on an estimated 0.2mm thick skull^130^ and a 1mm impact depth. For sham animals, the identical coordinates were used despite any real contusion, to serve as a negative control. Contusion voxels were excluded from all correlations. Since the two distances are spatially collinear (within-animal Spearman r ≈ 0.4–0.6), we isolated each predictor’s unique contribution with rank-based (Spearman) partial correlations. We report three quantities per group: the gradient–concordance partial, the anatomical–concordance partial, and their per-animal difference (gradient minus anatomical). For each, we tested whether it differed from zero within each group with a one-sample t-test after a Fisher z-transformation on the per-animal partial correlations. To ensure the gradient-concordance correlation observed in the CCI group was not artifactually inflated by larger connectivity changes in the CCI group compared to the shams, we performed voxelwise permutation (n=1000) of concordance values. If the gradient-concordance relationship was a real biological phenomenon, the random reshuffling would nullify the correlation. Fisher combination of p-values was performed on the correlation permutations to test if the true partial r significantly exceeded all 1,000 permuted values.

### dMRI analyses

dMRI scans were denoised, Gibbs-ringing corrected, and N4 bias corrected using *dwidenoise*, *mrdegibbs,* and *dwibiascorrect* implemented in *MRtrix3*^131^. Distortion and eddy current correction were then performed using the pairs of B0 images with opposite encoding directions via *MRtrix3*’s *dwifslpreproc* wrapper of *FSL*’s *topup* and *eddy*. The corrected volume was upsampled to 0.090mm isotropic voxel size, and the DSURQE atlas was registered to the T2-weighted structural scan. Finally, *dwi2tensor*, *tensor2metric*, and *AMICO*^132^ were used to obtain fractional anisotropy (FA), neurite density index (NDI), and orientation dispersion index (ODI) for each ROI in the atlas.

### Perfusion and fixation

After the final behavioral test, the mice were anesthetized with 5% isoflurane and perfused transcardially with cold 25ml phosphate buffered saline (PBS) containing 20U/ml heparin, followed by 25ml 4% paraformaldehyde (PFA) in PBS. The brains were fixed overnight in 4% PFA and subsequently stored in PBS.

### Tissue clearing, light-sheet fluorescence microscopy, and computational analysis

Whole mouse brains were processed with SHIELD to preserve tissue architecture, endogenous fluorescence and protein antigenicity^133^. The brains were then delipidated and immunolabeled via the Radiant protocol using the Smartbatch+ device (LifeCanvas Technologies). We used the following primary antibodies: rabbit host anti-c-Fos (3µg per brain, cat #ab214672, Abcam), rat host anti-GFAP (10µg per brain, cat #13-0300, ThermoFisher)); and secondary antibodies: c-Fos: Donkey anti-Rabbit IgG (H+L) Highly Cross-Adsorbed Secondary Antibody, Alexa Fluor™ 647 (6µg per brain, cat #A-31573, Invitrogen); GFAP: Donkey anti-Rat IgG (H+L) Highly Cross-Adsorbed Secondary Antibody, Alexa Fluor™ Plus 555 (20µg per brain, cat #A48270, Invitrogen)). Finally, samples were index-matched (n=1.52) by incubating in EasyIndex solution, and imaged with the Miltenyi UltraMicroscope Blaze at 4× magnification (EYFP, GFAP: 1.625µm x 1.625µm x 3µm; c-Fos: 1.625µm x 1.625µm x 4µm resolution).

Images were stitched with Imaris, converted to BIDS format via SPIMprep^134^, and registered to the adapted DSURQE mouse atlas via SPIMquant^135^. For segmentation of EYFP-expressing axons, we first applied contrast-limited adaptive histogram equalization enhancement before employing an in-house developed AI-based pipeline MAPL3 for axonal mapping^37^. Briefly, the 3D volume was divided into patches, preprocessed to correct for light-sheet artifacts, and segmented using a pre-trained 3D UNETR model with a binarization threshold of 0.2. Segmented patches were reassembled into a whole-brain volume and skeletonized via multi-threshold medial-axis transform with 15 erosion iterations. Skeletonization parameters (elongation threshold, linear mixing coefficient, and whether contrast-enhanced or unenhanced images were used for mixing) were optimized per sample. The output was voxelized to 40 µm isotropic resolution and region-level axon density was extracted based on the DSURQE atlas parcellations.

For c-FOS segmentation, we trained the AI-based Cartography of Ensembles (ACE) pipeline using 8 manually labelled (ground truth) image patches (112^3^ px with 1.625µm x 1.625µm x 4µm resolution). For GFAP segmentation, existing solutions yielded substandard predictions due to domain shift and artifacts. We thus trained our model using 8 manually labelled image patches (112^3^ px with isotropic 1.625µm^3^ resolution). Region-wise signal density and cell density were obtained for c-Fos+ neurons and GFAP+ astrocytes. Within-hemisphere normalization was performed for axon signal density, while whole-brain normalization was performed for GFAP signal density. Lateralization index was calculated as (right ROI normalized value − left ROI normalized value) / (right ROI normalized value + left ROI normalized value).

### Quantification and statistical analysis

Within-group behavioral readouts (chronic vs. baseline from the same subject) were analyzed using paired permutation t-tests (5000 permutations) via DABEST^136^. Cross-sectional groupwise differences (sham vs. CCI) across MRI and LSFM derived metrics were assessed using linear regression at each timepoint. To control for false discovery rates across *a priori* selected networks and ROIs, Benjamini–Hochberg corrections were applied, with uncorrected whole-brain analyses reported as exploratory. For metrics showing significant groupwise differences, we evaluated their relationship with memory performance via a linear regression. The change in metric readout value from baseline to chronic timepoint was correlated against the corresponding change in LD task accuracy. Due to the lack of baseline data for LSFM, the single chronic LSFM readout was regressed against the change in LD task accuracy. Statistical significance was assessed using Monte Carlo permutation testing (1000 permutations) to construct an empirical null distribution of the regression slope t-statistic^137^. Bias-corrected and accelerated (BCa) bootstrap^138^ resampling (1000 resamples) was applied to assess the stability of the R^2^ estimate, and the 95% confidence interval (CI) values are reported.

## SUPPLEMENTARY MATERIAL

**Supplementary Figure 1:**
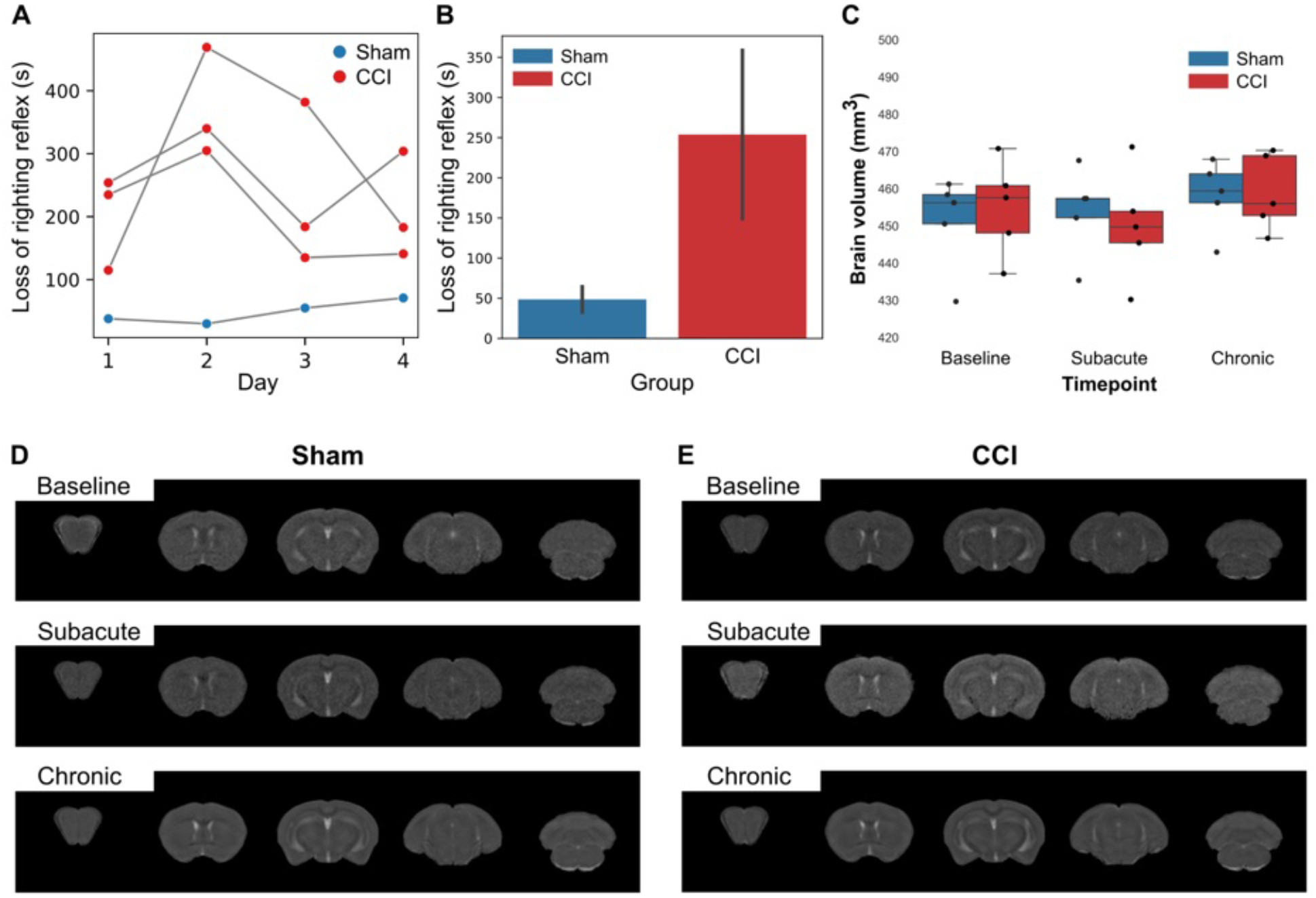
Gross sequelae of the CCI model. Characterisation of controlled cortical impact (CCI) severity (A) Duration of loss of righting reflex (LRR) of 1 sham and 3 CCI mice in CCI severity characterisation test on each day of hit. (B) Across all hits, CCI mice had a longer LRR (t-test, t = 3.66, *p <* 0.01). (C) Brain volume (mm3) was estimated from T2 structural scans, which showed no significant difference in brain volume between groups at any timepoints. Representative T2 structural scans of a sham (D) and a CCI (E) mouse, demonstrating absence of contrast.

**Supplementary Figure 2:**
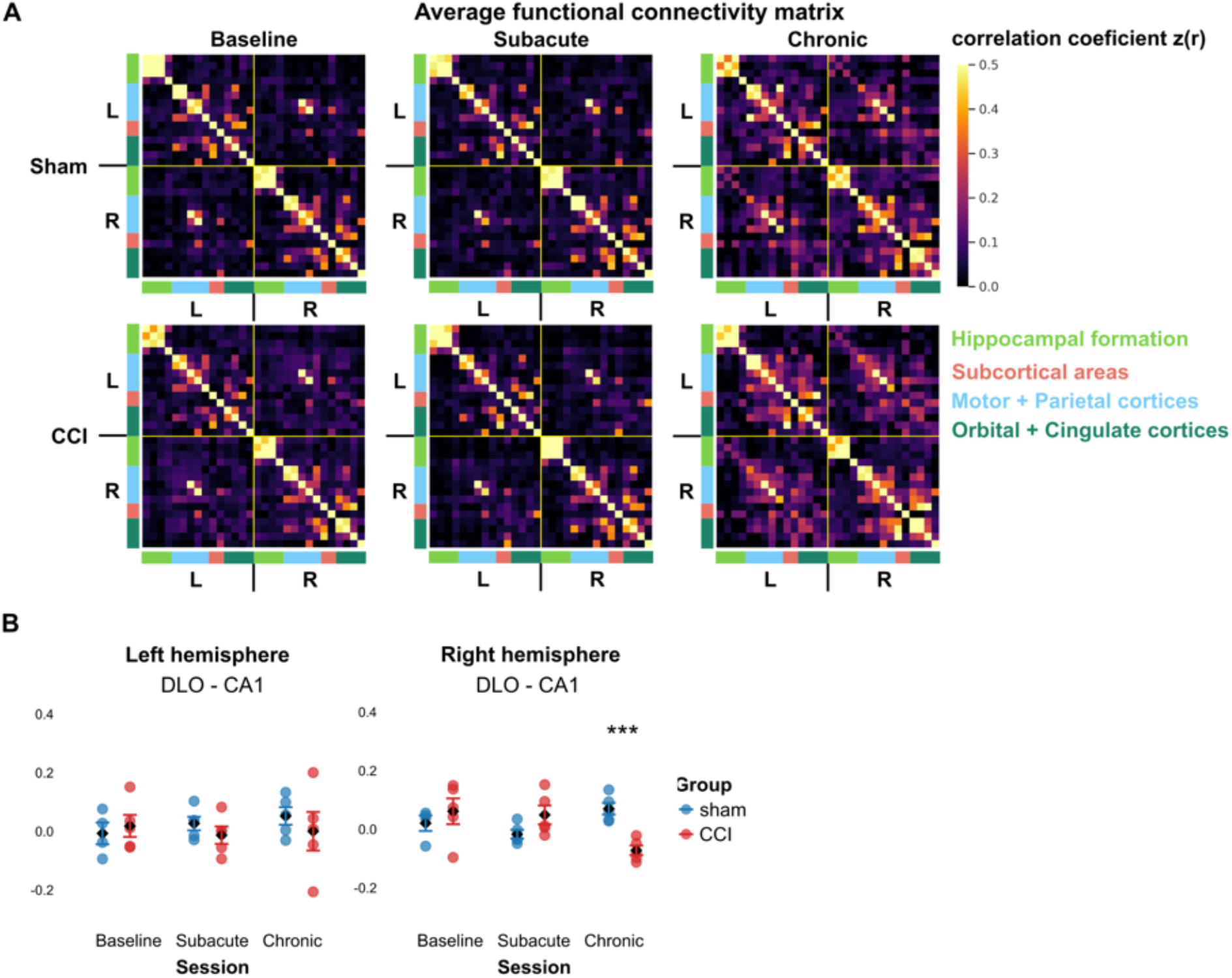
Resting-state fMRI results. (A) Groupwise averaged z-scored correlation heatmaps at each timepoint. Color shows group-averaged z-scored correlation. (B) ipsi-contusional DLO- CA1 connectivity was decreased significantly in the CCI group at chronic timepoint. Points represent individual animals colored by group, diamonds indicate group means and error bars represent standard error. CCI: controlled cortical impact; DLO: dorsolateral orbital cortex. Asterisks indicate *p*-value, where *p <* 0.05 *, *p <* 0.01 **, *p <* 0.001 ***.

**Supplementary Figure 3:**
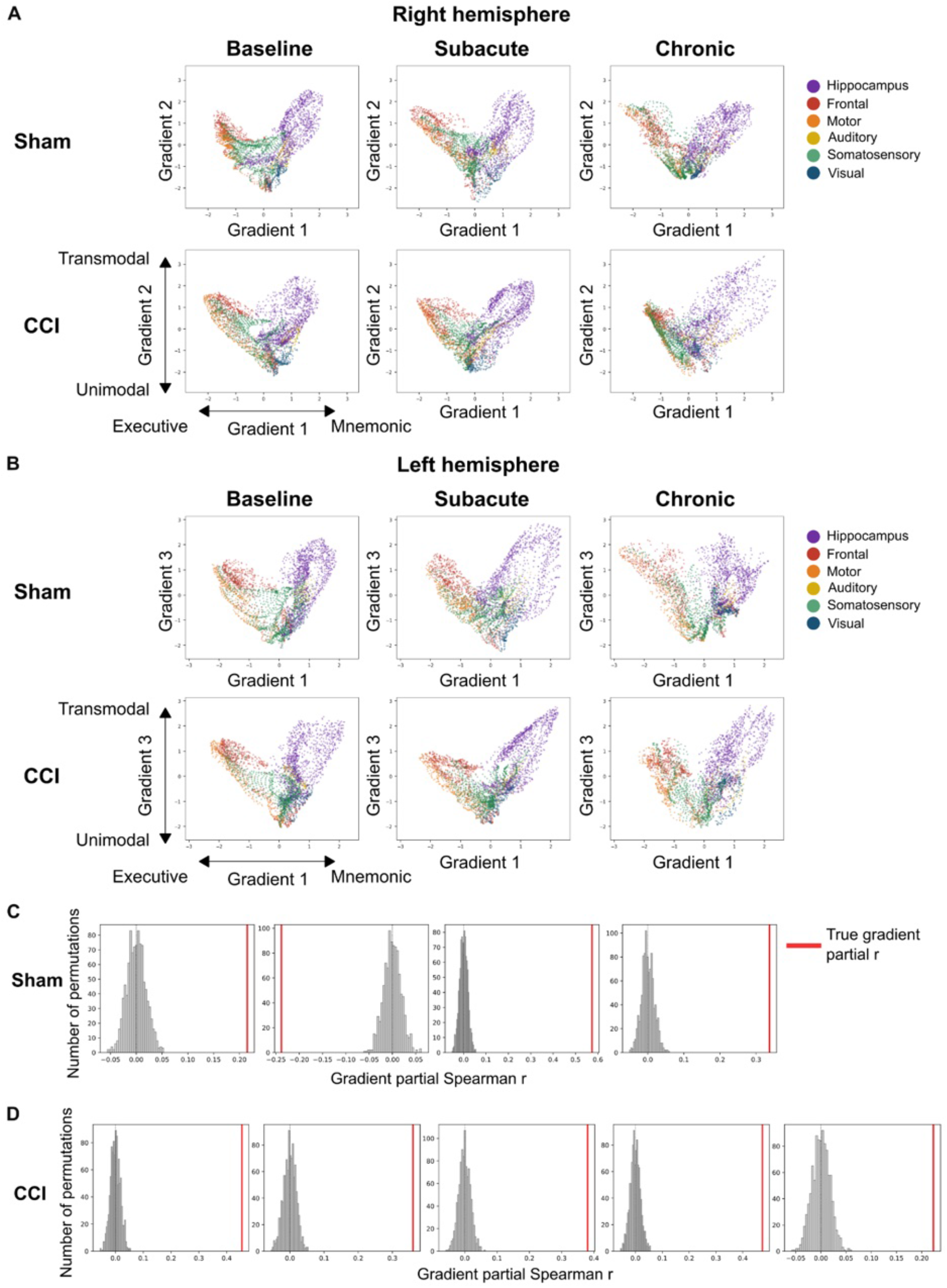
Functional connectomic gradient results. Group and timepoint averaged voxelwise gradients in gradient space defined by the executive - mnemonic and transmodal - unimodal gradients for the right ipsi-contusional (A) and left contra-contusional (B) hemispheres. Each point represents a voxel in the fMRI, colored by the functional network it belongs to. (C) Voxel-level permutation confirms spatial correspondence between gradient distance and concordance. For each animal, concordance values were randomly permuted across voxels 1,000 times and the gradient–concordance partial correlation recomputed on each permutation (grey histograms; null mean ≈ 0). Red lines mark the true partial r. Each plot represents a single animal. In three of four sham animals (shown in (C)), the true partial r exceeded all 1,000 permuted values (Fisher combined *p =* 1.7 × 10⁻⁶) while this was true for all CCI animals (shown in (D)) (Fisher combined *p =* 6.6 × 10⁻¹¹). This confirmed that the gradient– concordance relationship arose from the true spatial ordering of concordance values rather than an artifact of metric construction, and validated that the selective suppression of anatomical distance as a co-predictor of functional connectivity changes was a CCI-specific effect.

**Supplementary Figure 4:**
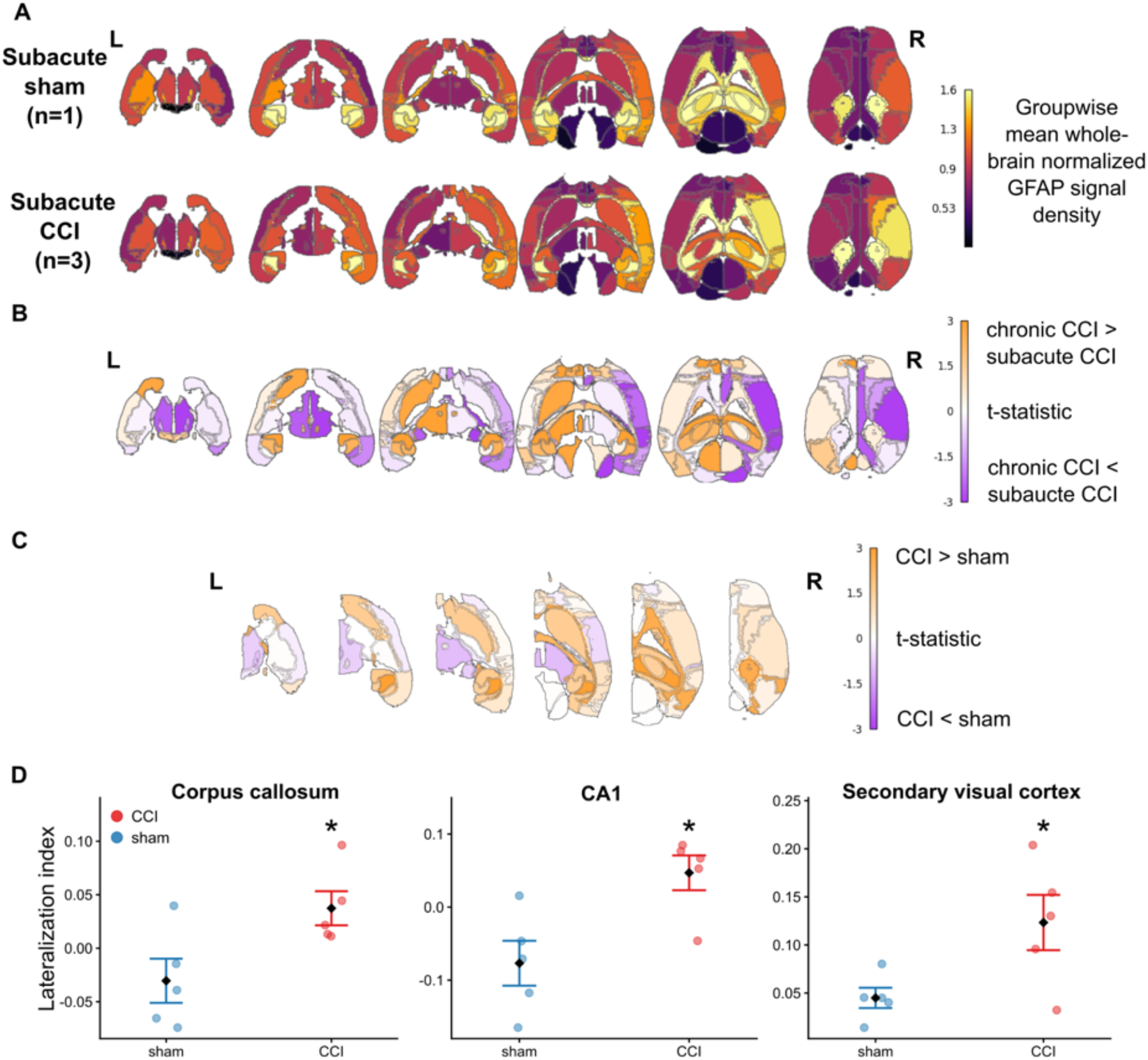
GFAP light-sheet fluorescence microscopy results. GFAP signal density elevation in subacute CCI, and increase of lateralization index of GFAP signal density in chronic CCI. (A) Group-averaged whole-brain normalized GFAP signal density maps show an increase in the subacute CCI group compared to the sham. Color scale shows group-averaged whole-brain normalized signal density from segmented GFAP+ astrocytes masks. (B) T-statistic from region-wise groupwise comparisons of LI. Color scale shows t-statistic. Regions in orange indicate higher whole-brain normalized GFAP signal in chronic CCI than subacute CCI while regions in purple indicate lower whole-brain normalized GFAP signal in chronic CCI than subacute CCI. (C) T-statistic from region-wise groupwise comparisons of LI at chronic stage. Color scale shows t-statistic. Regions in orange indicate higher LI in CCI than shams while regions in purple indicate lower LI in CCI than shams. (D) LI was increased in the corpus callosum, CA1 and secondary visual cortex in the CCI group. Points represent individual animals colored by group, diamonds indicate group means and error bars represent standard error. CCI: controlled cortical impact; LI: lateralization index. Asterisks indicate *p*-value, where *p <* 0.05 *, *p <* 0.01 **, *p <* 0.001 ***.

**Supplementary Table 1 -.** Region name abbreviations.

| Abbreviation | DSURQE atlas region name |
| --- | --- |
| CA1 | CA10r, LMol, CA1Rad, CA1Py |
| CA2/CA3 | CA2Py, CA20r, CA2Rad, CA3Py Inner, CA3Py Outer, CA30r, CA3Rad, SLu |
| DG | Dentate gyrus (MoDG, GrDG, PoDG) |
| EC | Entorhinal cortex (Caudomedial entorhinal cortex, Dorsal intermediate entorhinal cortex, Medial entorhinal cortex, Ventral intermediate entorhinal cortex) |
| DLO | Dorsolateral orbital cortex |
| LO | Lateral orbital cortex |
| MO | Medial orbital cortex |
| Cg | Cingulate cortex (areas 24a, 24a', 24b, 24b', 25, 29a, 29b, 29c, 30, 32) |
| FrA | Frontal association cortex |
| Str | Striatum (striatum, olfactory tubercle, fundus of striatum, nucleus accumbens) |
| Th | Thalamus |
| MOp | Primary motor cortex |
| SSp | Primary somatosensory cortex (barrel field, dysgranular zone, forelimb region, hindlimb region, jaw region, shoulder region, trunk region, upper lip region) |
| VISp | Primary visual cortex |
| mPPC | Medial parietal association cortex |

## Notes

### Competing Interest Statement

The authors have declared no competing interest.

